# Mitochondrial protein import stress augments α-synuclein aggregation and neurodegeneration independent of bioenergetics

**DOI:** 10.1101/2022.09.20.508793

**Authors:** Liam P. Coyne, Arnav Rana, Xiaowen Wang, Sanaea Bhagwagar, Yumiko Umino, Eduardo C. Solessio, Frank Middleton, Xin Jie Chen

## Abstract

Several genetic and environmental risk factors for Parkinson’s disease have been identified that converge on mitochondria as central elements in the disease process. However, the mechanisms by which mitochondrial dysfunction contributes to neurodegeneration remain incompletely understood. Non-bioenergetic pathways of the mitochondria are increasingly appreciated, but confounding bioenergetic defects are a major barrier to experimental validation. Here, we describe a novel bioenergetics-independent mechanism by which mild mitochondrial protein import stress augments neurodegeneration. We induced this mitochondrial protein import stress in an established mouse model of Parkinson’s disease expressing the A53T mutated form of α-synuclein (SNCA). Mice with import stress in addition to the A53T mutation demonstrated increased size of α-synuclein aggregates, co-aggregation of mitochondrial preproteins with α-synuclein, and worsened neurodegeneration. Importantly, we found no evidence of bioenergetic defects in any of the mutant mice, even with the added import stress. These data suggest that mitochondrial protein import stress contributes to neurodegeneration through cytosolic proteostatic stress and co-aggregation of mitochondrial and neuropathogenic proteins independent of bioenergetics. Given that protein import efficiency is affected by many types of mitochondrial stress, our findings add a new layer to understanding why the pathogenic mitochondrial dysfunction and cytosolic protein misfolding pathways converge in neurodegenerative diseases such as Parkinson’s disease.

## INTRODUCTION

Mitochondrial defects and cytosolic protein aggregation in neurons are co-manifested across neurodegenerative diseases such as Parkinson’s disease (PD), Lewy Body Dementia, Alzheimer’s disease, Amyotrophic Lateral Sclerosis (ALS) and Huntington’s disease. These two hallmarks are virtually ubiquitous despite significant variations in genetic and phenotypic makeup among diseases. While both hallmarks can individually drive neurodegeneration, whether and how they interact during initiation and/or progression of these complex diseases are poorly understood, and likely to be multifactorial. The effects of pathogenic cytosolic protein species on mitochondrial function have been extensively investigated ^1^. By comparison, whether and how mitochondrial function affects cytosolic proteostasis and pathological protein aggregation is vastly understudied.

Recent work in yeast and human cells has shown that a wide range of mitochondrial stressors can affect cell viability by reducing the import of mitochondrial preproteins causing their toxic accumulation and aggregation in the cytosol ^2–8^. This cell stress mechanism was termed mitochondrial Precursor Overaccumulation Stress (mPOS) ^9^. In the context of neurodegeneration, this raises the intriguing possibility that mitochondrial dysfunction could contribute to cytosolic protein aggregation, either by directly causing proteotoxicity or by amplifying preexisting proteostatic stress in the cytosol driven by misfolded pathogenic proteins (e.g., α-synuclein in PD) ^10^. The critical barrier to testing this idea has been lack of an adequate animal model of non-lethal mitochondrial protein import stress, especially one that isolates the effect of protein import defects in the absence of confounding bioenergetic defects. Such a model is essential for testing the pathogenicity of protein import stress and mPOS.

Modeling mitochondrial protein import stress in animals is particularly challenging because virtually all mitochondrial functions depend on efficient import through a limited number of protein translocase channels. All but 13 of the 1,000-1,500 mitochondrial proteins are encoded in the nucleus, synthesized by cytosolic ribosomes, and then imported into the organelle. The import machinery is intricate, relying on numerous chaperones both inside and outside mitochondria, as well as multiple pore-forming protein complexes in the inner and outer mitochondrial membranes ^11–13^. The sole entry point for >90% of mitochondrial proteins is the translocase of the outer membrane (TOM) complex. We previously engineered mutant variants of the mitochondrial ADP/ATP carrier protein that clog the TOM complex to partially obstruct general protein import ^14^. The best-characterized clogger mutant protein was Ant1^p.A114P,A123D^, encoded by *Slc25a4 ^p.A114P,A123D^*, which we used to generate a knock-in mouse model of protein import stress. In the *Slc25a4 ^p.A114P,A123D^* /+ mice (or “clogger” mice), unimported mitochondrial proteins accumulate in the cytosol of skeletal muscle, which correlates with an age-dependent mitochondrial myopathy. Interestingly, approximately 4% of the mice become paralyzed around 12 months of age, suggesting a direct role of protein import clogging in neurodegeneration. However, the vast majority (∼96%) of clogger mice have a normal lifespan, and do not develop overt neurological symptoms. This observation suggests that mitochondrial protein import is only mildly clogged. In the present study, we utilize these mice to study the specific effects of moderate protein import stress on neurological function. Most importantly, we then used mouse genetics to test whether and how protein import stress might contribute to neurodegeneration in a mouse model of PD that involves the cytosolic misfolding and aggregation of a mutant form of α-synuclein (A53T), that is a dominant cause of familial PD.

## RESULTS

### Ant1^p.A114P, A123D^ does not affect mitochondrial bioenergetics in the brain and does not impair motor coordination or cognitive function

The human ANT1^p.A114P^ protein is pathogenic, causing autosomal dominant Progressive External Ophthalmoplegia (adPEO) ^15^. In addition to PEO and skeletal muscle phenotypes, ANT1-induced adPEO also associates with neurological and psychiatric phenotypes such as sensorineural hearing loss, cognitive impairment, dementia, bipolar and schizoaffective disorders ^59–63^. ANT1^p.A123D^ is associated with myopathy and cardiomyopathy ^16^. The Ant1^P.A114P,A123D^ /+ “clogger” mice were generated to facilitate phenotypic scoring and evaluate the *in vivo* effects of protein import clogging, as two mutations together resulted in direct TOM complex clogging ^14^. To evaluate mitochondrial bioenergetics in the central nervous system of non-paralytic clogger mice, we measured oxygen consumption from purified brain mitochondria. Unlike skeletal muscle ^14^, brain mitochondria from clogger mice at the age of 9 and 24 months do not have reduced maximal respiratory rate (state 3) when stimulating complex I-based respiration (Figure 1A-B). ADP-depleted respiration (state 4) and the respiratory control ratio (state 3/state 4) were also unchanged (Figure 1C-D). These observations suggest that mitochondrial oxygen consumption is well-coupled with ATP synthesis and the maximal complex I-based respiratory capacity is not reduced in clogger brain mitochondria. Mitochondrial respiration was also unaffected when complex I is inhibited with rotenone and complex II is stimulated with succinate (Figure 1E-H). Thus, mitochondrial respiration is unaffected or minimally affected in mouse brain. Consistent with this, we assessed the steady-state levels of respiratory chain proteins by immunoblot and found that representative subunits of complexes I, II, III and V were unaffected in clogger brain mitochondria (Figure 1I). We extended our analysis to the spinal cord by directly examining mitochondrial morphology using transmission electron microscopy. The data showed preserved mitochondrial morphology in ventral horn neurons in the non-paralytic clogger mouse spinal cord (Figure 1J). Taken together, these data suggest that mitochondrial structure and bioenergetic function are generally preserved in the central nervous system of clogger mice.

**Figure 1.**
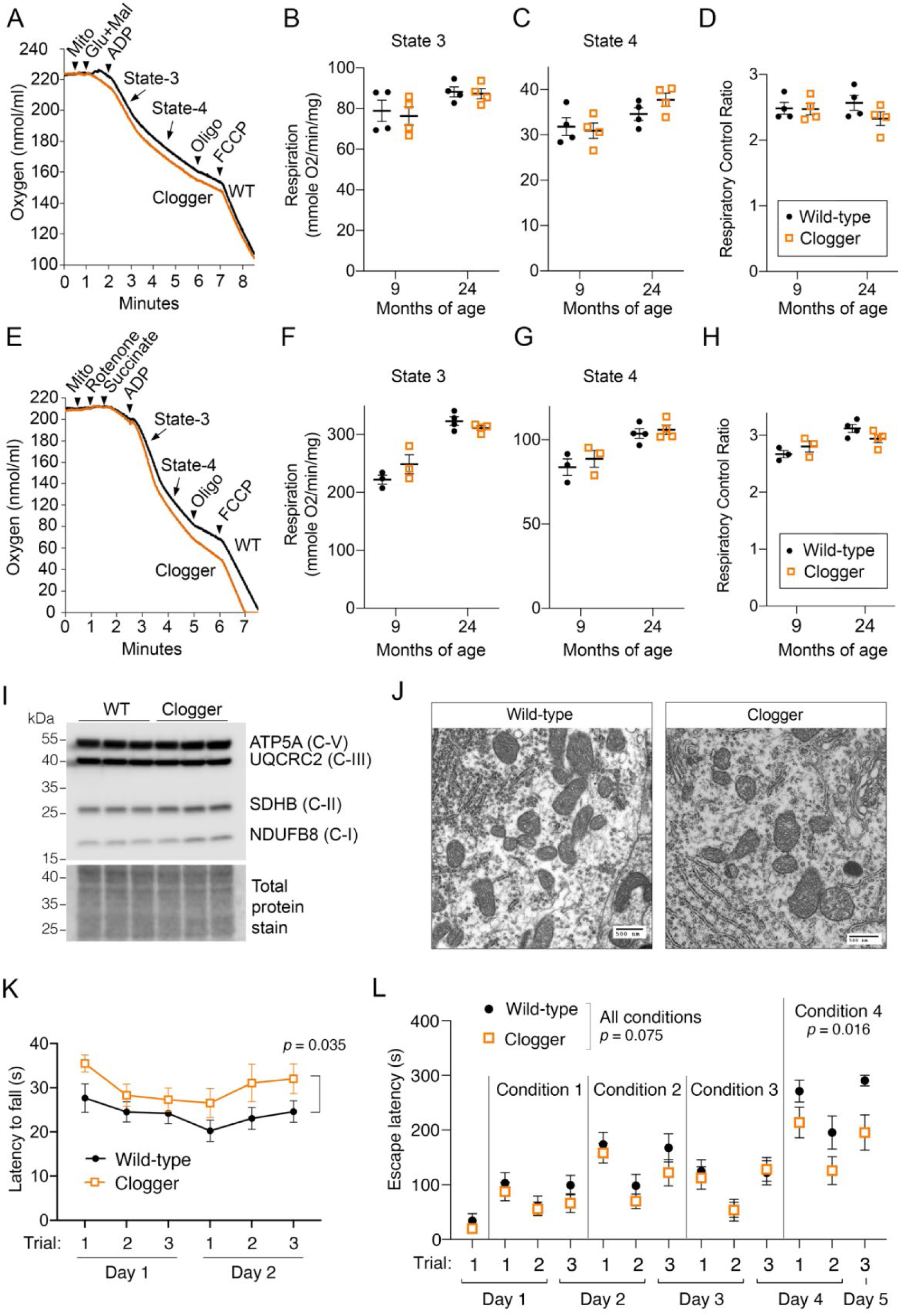
Mild mitochondrial protein import clogging with Ant1^p.A114P,A123D^ does not affect mitochondrial bioenergetics but moderately improves motor coordination and executive function. (A) – (D) Respirometry of purified brain mitochondria with complex I stimulated by glutamate (glu) and malate (mal). N = 4 mice/genotype/age group. Two measurements were taken per mouse, the average of which is plotted in (B)-(D). Data from different ages were analyzed separately due to a suspected batch effect. Independent repeated measures ANOVA within each age group (with measurement order as the within-subjects variable and genotype as the between-subjects variable) confirmed no effect of measurement order or genotype for any measure potted in (B)-(D) or (F)-(H). FCCP, Trifluoromethoxy carbonylcyanide phenylhydrazone; Oligo, oligomycin. State 3 respiration is the maximal respiratory rate after addition of ADP. State 4 respiration is the respiratory rate after depletion of ADP. The respiratory control ratio is State 3 divided by State 4 respiratory rates. Decreased respiratory control ratio would be interpreted as increased mitochondrial damage. Error bars in these and all subsequent panels and figures indicate standard error of the mean (SEM). (E) – (H) Respirometry of purified brain mitochondria with complex II stimulated by succinate and complex I inhibited by rotenone. N = 3 mice/genotype at 9 months of age and 4 mice/genotype at 24 months of age. Data collected and analyzed as in (A)-(D). (I) Immunoblot analysis of representative subunits of the respiratory complexes from purified brain mitochondria. (J) Transmission electron microscopic analysis in the cell body of spinal cord ventral horn neurons showing no obvious changes to mitochondrial ultrastructure in non-paralytic clogger mice. Scale bar is 500 nm. (K) Clogger mice perform better than wild-type on the accelerating rotarod at 19-22 months of age, n > 16 mice per genotype, n > 5 per sex per genotype. Data were analyzed with a two-way repeated measures ANOVA probing for an effect of sex and genotype, with Geisser-Greenhouse correction. Depicted is the main effect of genotype. (L) Clogger mice have moderately enhanced executive function, as judged by speed in removing obstacles from a 5 cm x 5 cm doorway to escape from a stressful environment. Mice were 20-23.5 months of age; n = 13 wild-type (5 females, 8 males), 15 cloggers (9 females, 6 males). Different “Conditions” are different obstacles. Data analyzed by repeated measures ANOVA with Geisser-Greenhouse correction.

To explore the effects of mitochondrial protein import clogging on neurological function, we performed an array of behavioral assays on non-paralytic clogger mice. To ensure validity of the behavioral assays, we first assessed basic vision and motor function. Using optomotor response testing, we confirmed that visual acuity and contrast sensitivity are preserved in clogger mice (Figure S1A-B). While 30-month-old clogger mice clearly exhibit skeletal muscle atrophy and weakness ^14^, this effect appears to be age-dependent, as mice at the age of ∼13 months have preserved muscle function, as suggested by treadmill exhaustion testing (Figure S1C-D) and swim speed (Figure S1E). We also tested ∼20-month-old mice on an accelerating rotarod to measure motor coordination and balance. Surprisingly, we found that the clogger mice performed significantly better than wild-type mice (Figure 1K). This curious finding has previously been observed in another mouse model of mitochondrial disease ^17^.

To assess general locomotor activity, we performed an open field test. When left undisturbed in a brightly lit open field for 10 minutes, female clogger mice move at a faster average speed compared with wildtype (Figure S2A), reminiscent of another mouse model of mitochondria-induced Progressive External Ophthalmoplegia ^18^. However, increased locomotor activity could not be attributed to increased anxiety-like behavior (Figure S2B-C). To assess cognitive function, we monitored animal behavior in several assays sensitive to short-term and long-term memory as well as executive function. The Y-maze spontaneous alternation test relies on rodents’ natural tendency to explore new areas. When they repeatedly return to arms of the Y-maze that they most recently visited, this is interpreted as a defect in working spatial memory. Results from this assay suggested that clogger mice have preserved spatial working memory (Figure S2D). The novel object recognition test relies on rodents’ natural tendency to explore new objects after becoming familiarized with other objects over multiple prior training days. The data from this assay suggested that clogger mice have preserved long-term recognition memory (Figure S2E). The Morris water maze tests rodents’ ability to learn and remember the location of a hidden escape platform that they must swim to, which is unpleasant for rodents. Results from this assay suggested that clogger mice have preserved long-term spatial memory and learning (Figure S2F-H). Collectively, these data suggest that learning and both short-term and long-term memory are intact in clogger mice.

Finally, we tested executive function, which refers to higher-level processing used to control and coordinate behavior in response to changing task demands. To test this, we used a “puzzle box” assay in which mice are placed in a stressful environment and the amount of time it takes to escape through an obstructed exit doorway is used as a proxy for executive function ^19, 20^. We found that, for the most difficult obstruction (“Condition 4”), clogger mice were significantly better at escaping the stressful environment (Figure 1L). Therefore, clogger mice may have improved executive function in a manner that cannot be attributed to increased anxiety-like behavior.

Overall, these behavioral data are critical for establishing that Ant1^p.A114P,A123D^ alone does not cause neurological dysfunction in non-paralytic mice. In fact, motor coordination and executive function appear to be equal or better than in wildtype mice.

### Transcriptional remodeling in the central nervous system

To learn whether Ant1^p.A114P, A123D^-induced protein import clogging triggers specific stress responses, we first analyzed the spinal cord transcriptome because the spinal cord appears to be preferentially affected in clogger mice ^14^. We found a robust transcriptional signature of 149 differentially expressed genes (*q* < 0.05) (Figure 2A, Supplementary Table 1). Among the four most upregulated genes were *Hspa1b*, *Igf2* and *Igfbp6* (Figure 2B). *Hspa1b* encodes one of the major stress inducible Hsp70 chaperones, which are important for protein folding and protection against proteostatic stress. Hsp70 is also involved in the stabilization and mitochondrial delivery of mitochondrial preproteins in the cytosol ^21^. HSP70 genes are also transcriptionally activated by mitochondrial protein import clogging in human cells ^5^ and yeast ^14, 22^. These observations suggest that transcriptional upregulation of HSP70 genes is a conserved response to mitochondrial protein import clogging.

**Figure 2.**
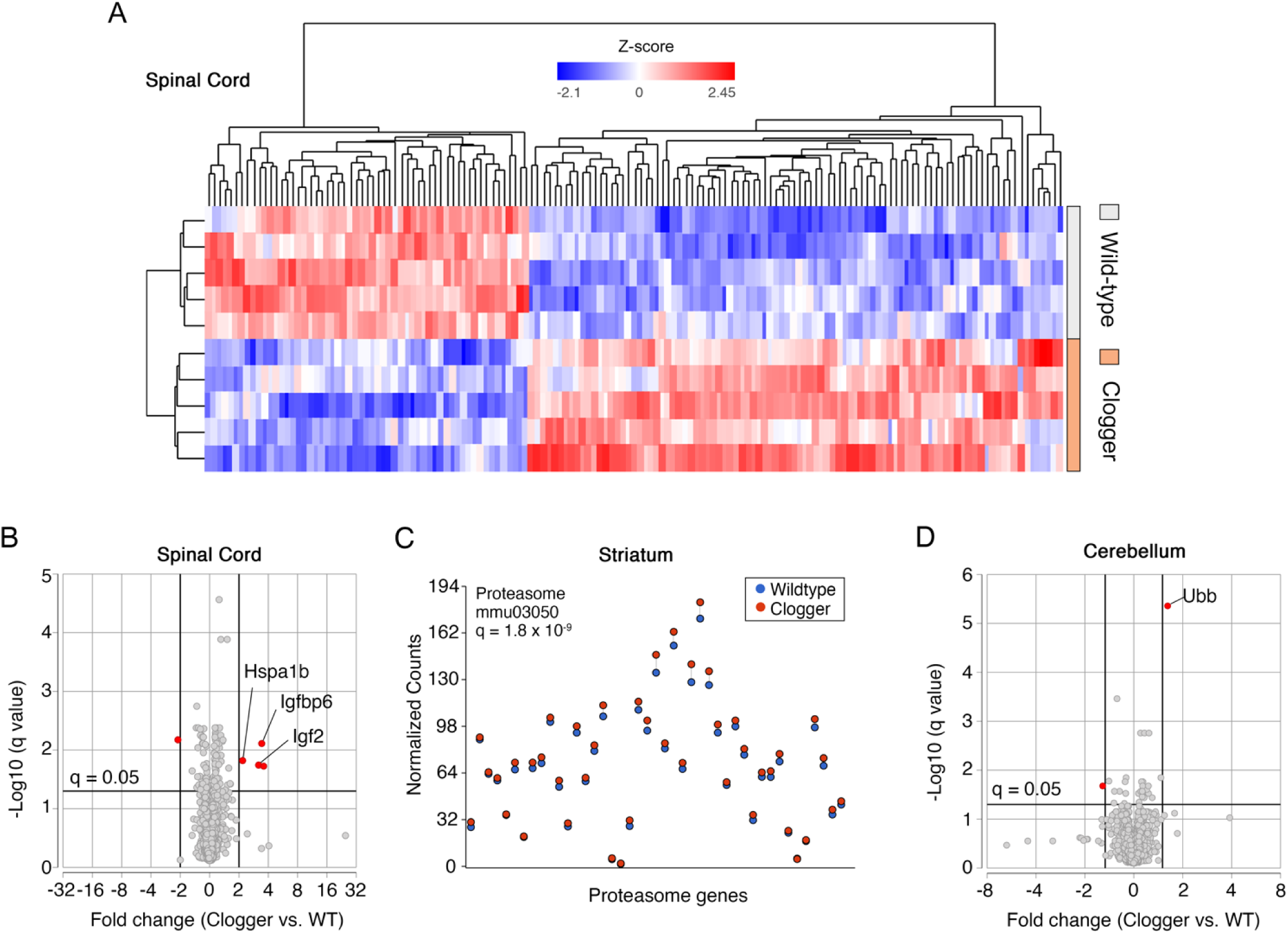
Transcriptomic analysis of neural tissues suggests cytosolic proteostatic stress in the clogger mice. (A) Heatmap of all genes that are differentially expressed in the spinal cord of clogger mice, compared with wild-type (*q*<0.05). (B) Volcano plot of transcriptomics from the spinal cord at 30 months of age. (C) Partek Pathway analysis of transcriptomics data from the striatum at 24 months of age suggests global proteasome upregulation. (D) Volcano plot of transcriptomics from the cerebellum at 30 months of age shows significant upregulation of *Ubb*. Note that *Slc25a4* (*ANT1*) transcript level is consistently lower in the spinal cord (C) and cerebellum (D) from the Clogger mice due to partial instability and/or reduced transcription from the mutant *Ant1^p.A114P, A123D^* allele (our unpublished observation).

Insulin-like growth factor-binding proteins (IGFBPs) are secreted proteins that bind insulin-like growth factors (IGFs) to regulate their transportation, localization, and function. IGFBP-6 is unique to this family in that it has a 20- to 100-fold higher affinity for IGF2 compared with IGF1 ^23^. Thus, co-induction of *Igfbp6* and *Igf2* in clogger-mouse spinal cords may be a coordinated stress response. Recent studies demonstrated that IGF2 can protect against neurodegeneration through extracellular disposal of protein aggregates ^24–26^.

We extended our RNA-Seq analysis to other regions within the central nervous system. In contrast to the spinal cord, we observed much more restricted transcriptional changes at the individual gene level in the cerebellum and striatum, with only 28 and 6 differentially expressed genes respectively (*q* < 0.05) (Figure S3, Supplementary Tables 2-3). Notably absent were any changes in genes involved in oxidative phosphorylation, consistent with a lack of mitochondrial respiratory defects. Instead, pathway analysis revealed global upregulation of proteasomal genes in the striatum (Figure 2C), which would be predicted if protein import clogging is causing mitochondrial Precursor Overaccumulation Stress (mPOS) in the cytosol ^9, 10, 22, 27^. Consistent with disturbed cytosolic proteostasis, we found that *Ubb* (ubiquitin B) is the most upregulated gene in the cerebellum (Figure 2D). *UBB* transcription was previously shown to be upregulated in neurons of Parkinson’s disease patients ^28^. Related to the *Ubb* findings, we also noted significant increases in the *Dnajb11* transcript, which encodes an Hsp40 chaperone involved in unfolded protein responses and proteostasis (Supplementary Tables 2). Finally, we point out that there was moderately but significantly increased expression of *Comt* transcript detected in the clogger cerebellum as well, which is a gene involved in the degradation of dopamine and norepinephrine that is commonly targeted by drugs used to treat patients in the early stages of PD. Overall, these transcriptional findings are consistent with extra-mitochondrial proteostasis disruption by mitochondrial protein import clogging. Moreover, different regions of the central nervous system seem to have different stress responses.

### Protein import clogging worsens neurodegenerative phenotype in a mouse model of Parkinson’s disease

We next used mouse genetics to test whether mild mitochondrial protein import stress with Ant1^p.A114P,A123D^ can worsen cytosolic protein aggregation and disease in an established mouse model of neurodegeneration. We crossed the clogger mice with hemizygous transgenic mice expressing the human α-synuclein with the A53T mutation, which is an autosomal dominant cause of familial Parkinson’s disease and a highly aggregation-prone protein ^29^. Transgenic α-synuclein(A53T) mice (hereafter referred to as “α-syn” mice) develop overt neurological dysfunction at 12-16 months of age, which progresses to end-stage paralysis within 2-3 weeks of symptom onset ^29^. Thus, to detect any additive effect of Ant1^p.A114P,A123D^ expression, we characterized the mice at 8-9 months of age (∼20 mice per sex per genotype).

Motor symptoms such as bradykinesia and postural instability are a hallmark of PD. We first assessed balance and coordination using the beam walking test. Mice were trained to walk across a thin beam and the number of times their paws slipped off the beam was scored as a measure of coordination (Figure 3A). As expected, α-syn mice slipped more often on both a 0.5-inch diameter cylindrical beam (*p* = 1.3 x 10^-5^), as well as a 0.25-inch rectangular beam (*p* = 7.2 x 10^-6^) Figure 3B-C). Ant1^p.A114P,A123D^ expression drastically enhanced this phenotype on the smaller beam (*p* = 3.9 x 10^-6^), while showing a trend for further impairing performance on the larger beam. Importantly, Ant1^p.A114P,A123D^ expression had no effect in the wild-type background (Figure 3B-C). This suggests that mitochondrial protein import clogging and α-synuclein(A53T) expression act synergistically to impair motor function.

**Figure 3.**
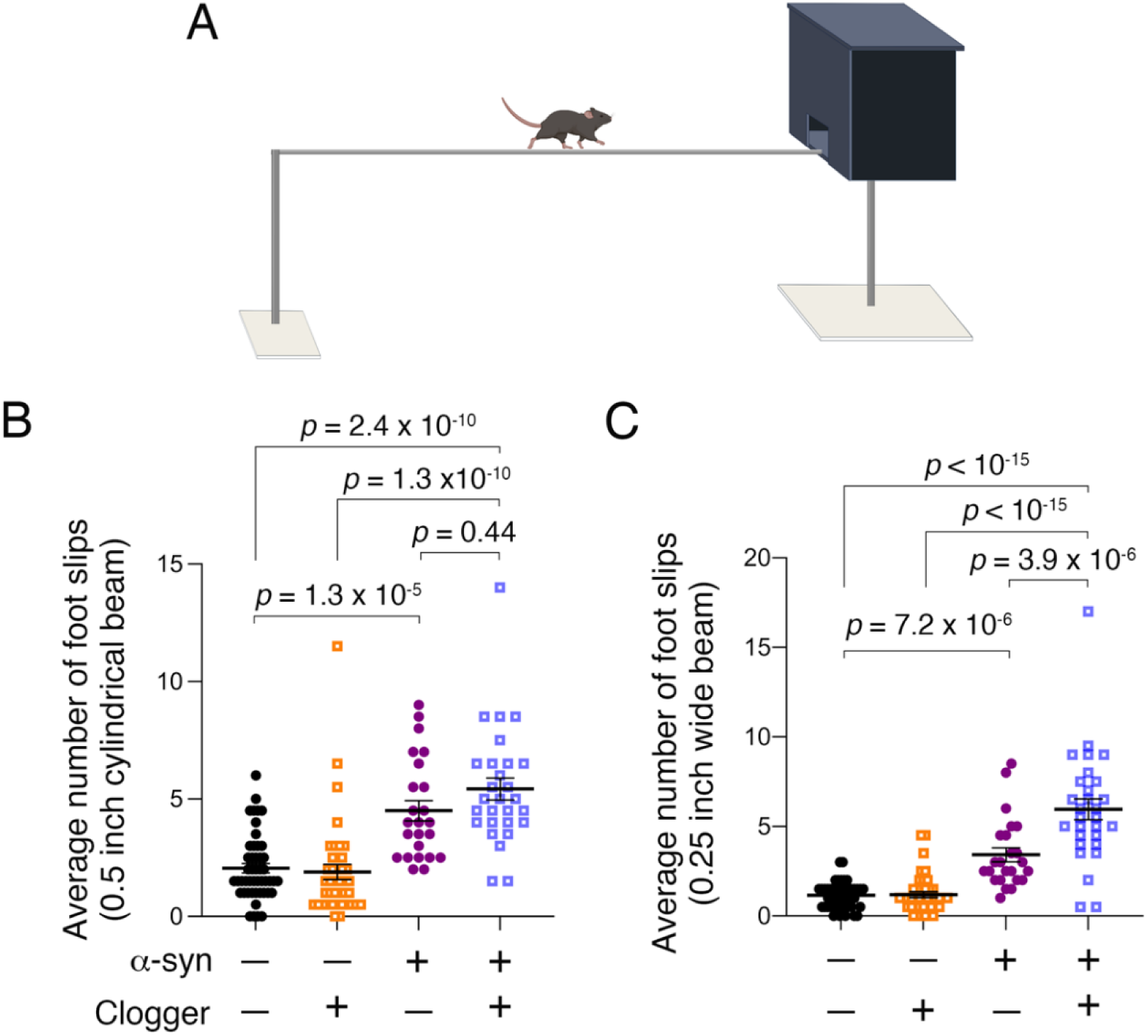
Mild protein import stress impairs motor coordination only in the α-synuclein(A53T) transgenic background. (A) Schematic of the beam walking test. (B) Beam-walking test suggested mildly worsened coordination on a 0.5-inch cylindrical beam in Clogger + α-Syn A53T double mutant compared with α-Syn A53T single mutant mice, as judged by the number of times each mouse’s hind limbs slipped off the beam. Data were first analyzed with a three-way ANOVA probing for effects of sex, α-synuclein, and clogger genotype. With no significant effect or interaction effect from sex, males and females were consolidated. Adjusted *p*-value that is shown is from a two-way ANOVA with Sidak’s multiple comparison’s test. (C) Beam-walking test suggested severely impaired coordination on a 0.25-inch rectangular beam in Clogger + α-Syn A53T double mutant compared with α-Syn A53T single mutant mice. Data analyzed as in (B). There was a significant interaction between α-synuclein and clogger genotype (*p* = 7.5 x 10^-5^).

We followed these mice through end-stage paralysis to determine if Ant1^p.A114P,A123D^ expression reduced lifespan in α-syn mice. We found no change in median lifespan in double mutant mice compared with α-syn alone (Figure S4A). Intriguingly, we noticed that in the longest-lived mice (upper quartile), Ant1^p.A114P,A123D^ expression significantly reduced maximum lifespan (log-rank *p* = 0.04 (Figure S4A), *t* test *p* = 0.028 (Figure S4B)). In sum, the data show that protein import clogging by Ant1^p.A114P,A123D^ can worsen motor deficits and modestly shorten maximum lifespan in a mouse model of α-synuclein-induced PD.

Does protein import clogging aggravate additional phenotypes of α-syn mice? Up to 40% of PD patients experience anxiety ^30^. We found that mutant α-syn expression significantly increased anxiety-like behavior in the open field test (*p* = 1.3 x 10^-4^) (Figure S5A). While in the center zone, double mutant mice remained closest to the edge zone on average, suggesting Ant1^p.A114P,A123D^ expression moderately increases anxiety-like behavior in α-syn, but not wild-type mice (Figure S5B). Spontaneous locomotor activity was increased in α-syn mice (*p* < 10^-15^), as previously reported ^31^, which was not affected by Ant1^p.A114P,A123D^ (Figure S5C). In the novel object recognition test, α-syn expression caused a significant defect in long-term object memory (*p* = 2 x 10^-6^), with no effect from Ant1^p.A114P,A123D^ expression (Figure S5D). The total time spent exploring either object was largely unaffected by genotype (Figure S5E). In the Y-maze spontaneous alternation test, α-syn expression caused a significant defect in short term spatial memory (*p* = 2.5 x 10^-5^), with no effect of Ant1^p.A114P,A123D^ expression (Figure S5F). The total number of arm entries in the Y-maze was increased by α-syn expression (*p* = 1.6 x 10^-4^), with no effect of Ant1^p.A114P,A123D^ expression (Figure S5G). It is likely that the memory defects in α-syn mice are already severe by 9 months old, precluding detection of any potential additive effect from Ant1^p.A114P,A123D^ expression.

### Ant1^p.A114P, A123D^ does not affect mitochondrial bioenergetics in α-syn mice

There are substantial data to suggest that α-synuclein and other protein aggregates can impact mitochondrial bioenergetics, while other studies failed to generate consistent results ^1^. We tested the possibility that a synergistic defect in mitochondrial respiratory function may contribute to the enhanced double mutant phenotypes. We again measured respiration from complex I- and complex II-stimulated purified brain mitochondria. We found that α-synuclein(A53T) expression did not reduce respiratory rates in the tissues we examined at 9 months of age. More importantly, no respiratory deficiency was observed even in the Ant1^p.A114P,A123D^ plus α-syn double mutant mice (Figure 4A-H). Because Percoll-purified mitochondria are mostly non-synaptosomal, and α-syn primarily localizes to pre-synaptic sites, we also tested respiration from the synaptosomal brain fractions. These fractions are impure, containing microsomal membranes and myelin. Nonetheless, respiratory rates from synaptosomal fractions suggested that wild-type, A53T α-syn, and double mutant mice all have similar respiratory rates (Figure S6A-B). These data suggest that mitochondrial bioenergetic defects are not the cause of motor coordination defects in the α-syn single or Ant1^p.A114P,A123D^ plus α-syn double mutant mice.

**Figure 4.**
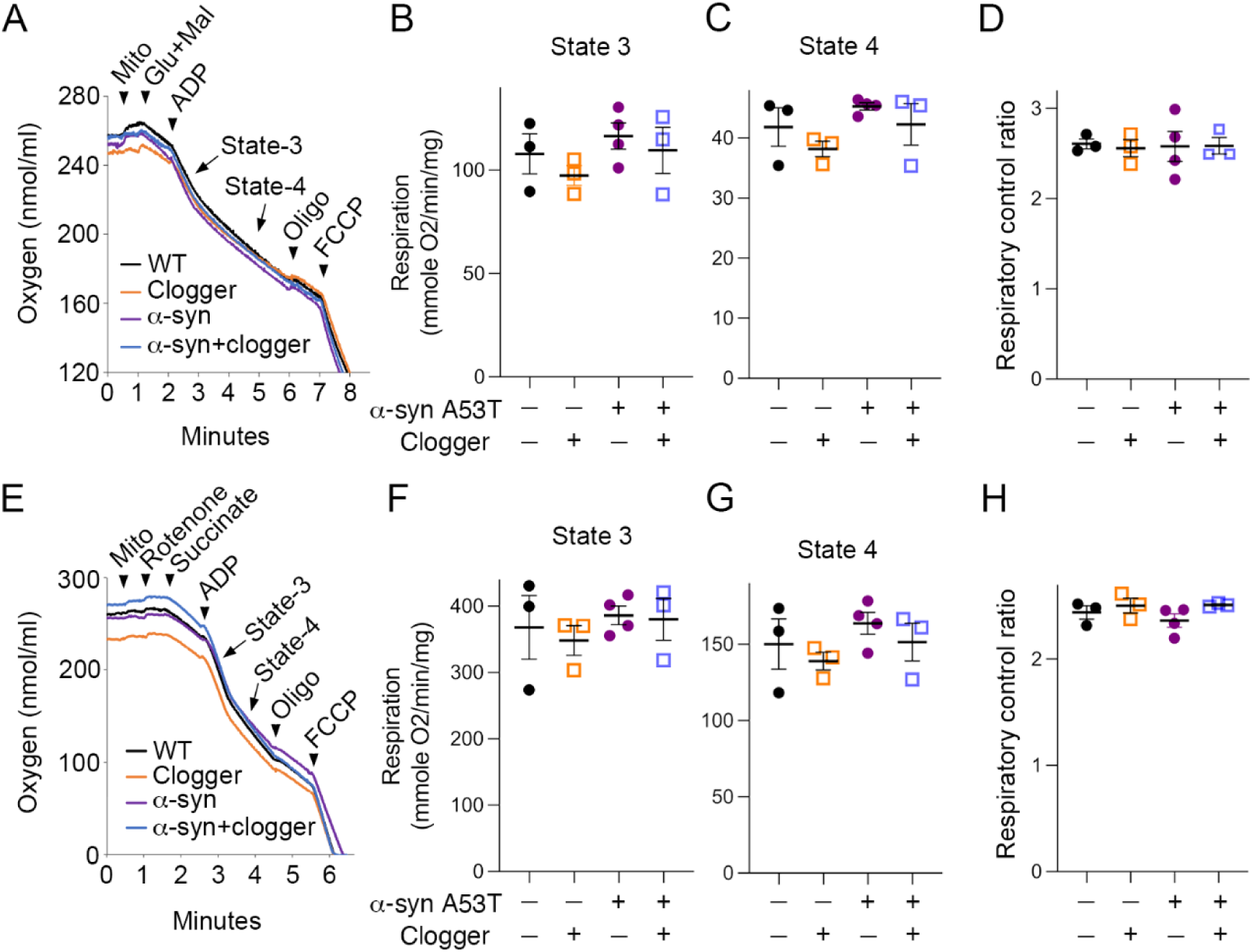
There is no detectable respiratory deficit in 9-month-old α-syn or α-syn plus clogger double mutant mice. (A) – (D) Respirometry of purified brain mitochondria with complex I stimulated by glutamate (glu) and malate (mal). 2 measurements were taken per mouse, the average of which is shown as a data point in (B)-(D). FCCP, Trifluoromethoxy carbonylcyanide phenylhydrazone; Oligo, oligomycin; Glu, glutamate; Mal, malate. State 3 respiration is the maximal respiratory rate after addition of ADP. State 4 respiration is the respiratory rate after depletion of ADP. The respiratory control ratio is State 3 divided by State 4 respiratory rates. Decreased respiratory control ratio would be interpreted as increased mitochondrial damage. Two measurements were performed per mouse. Each dot represents the average value from each mouse. (E) – (H) Respirometry of purified brain mitochondria with complex II stimulated by succinate (glu) and complex I inhibited by rotenone. Each dot represents the average value from each mouse.

Previous work showed that Cytochrome *c* oxidase (COX) activity was reduced in the spinal cord of end-stage α-syn mice, raising the possibility that this could be driving our motor coordination phenotype in non-paralyzed mice ^32^. However, we found that COX activity in spinal cord mitochondria from α-syn mice was preserved at 9-months old, the age at which we observe motor defects. Double mutant mice also showed no reduction in COX activity (Figure S6C-D). Taken together, these data strongly indicate that bioenergetic defects do not underlie the synergistic effect between mitochondrial protein import clogging and α-syn proteotoxicity that aggravates neurodegeneration.

### Mitochondrial protein import clogging increases the size of phosphorylated α-synuclein aggregates

We hypothesized that mitochondrial protein import clogging would cause mPOS and increase proteostatic burden in the cytosol, which may in turn increase α-synuclein aggregation. To narrow our focus to a particular region of the central nervous system, we performed detergent solubility studies to assess where α-synuclein is preferentially detergent-insoluble, which we use as a proxy for aggregation. We probed the Triton X-100-insoluble fractions from the forebrain, midbrain, cerebellum and spinal cords of end-stage α-syn mice for α-synuclein phosphorylated at serine 129 (P-α-syn). P-α-syn accumulates in pathological aggregates and is a more specific marker for protein aggregation and disease activity compared with non-phosphorylated α-synuclein ^33^. We found that insoluble P-α-synuclein preferentially accumulates in the spinal cord (Figure S7), consistent with previous studies ^29^. We also note that paralytic ^14^ and non-paralytic clogger mice have preferential effects on the spinal cord. The latter conclusion is based on the motor phenotype and correlative activation of more drastic transcriptional stress response pathways. We therefore focused our studies of protein aggregation on the spinal cord.

To quantitatively assess the size and number of α-synuclein aggregates, we performed immunofluorescence on the lumbar spinal cord. At 7 months of age, neither α-syn nor double mutant mice showed significant pathology when probing for P-α-syn (Figure 5A). In end-stage mice, we found many puncta staining positive for P-α-syn, suggesting widespread protein aggregation as expected (Figure 5A). We found that the average size of P-α-syn puncta was increased in the ventral horns of double mutant mice compared with α-syn only (Figure 5B-D). Consistent with increased puncta size, the average perimeter of puncta was also increased in double mutants (Figure 5E). We also observed an increased integrated density (and therefore larger size) of puncta in the double mutants (Figure 5F), despite the total area of P-α-syn in the double mutants being reduced (Figure 5G). These data support an overall increased propensity of α-syn aggregation in the double mutant relative to α-syn single mutant mice.

**Figure 5.**
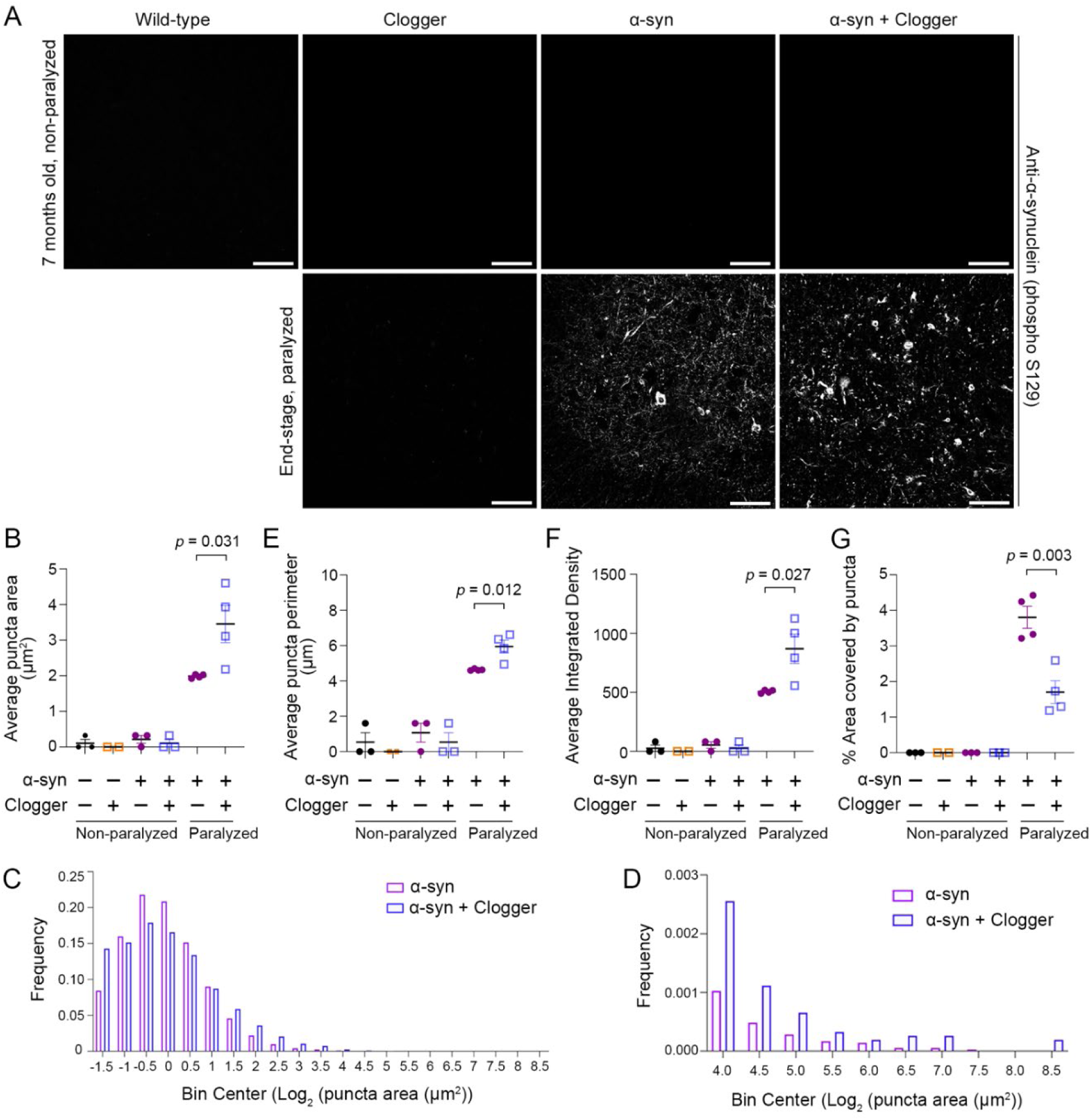
Increased size and intensity of α-syn phospho-S129 puncta in double mutant mice. (A) Representative images from immunofluorescence for α-synuclein phosphorylated at S129 in the ventral horns of the lumbar spinal cord from control (Wild-type, Clogger) and end-stage α-syn mice with or without protein import clogging. Scale bar = 100 microns. (B) Increased average P-α-syn puncta area in double mutant mice. (C) Frequency distribution of P-α-syn puncta size. (D) Frequency distribution as in (C), but focused on larger puncta that are not visible in (C). (E) Increased average P-α-syn puncta perimeter in double mutant mice. (F) Increased average integrated density of P-α-syn puncta in double mutant mice. (G) Reduced % area covered by P-α-syn puncta in double mutant mice. Aggregate data were generated exclusively from the ventral horns of three sections per mouse at anatomically distinct lumbar spinal cord levels from four biological replicates per genotype. *P* values were derived from student’s *t* test.

### Protein import clogging increases co-aggregation of mitochondrial proteins with α-synuclein

If cytosolic proteostatic stress from un-imported mitochondrial preproteins is increasing α-syn aggregation, then we would expect increased mitochondrial preprotein co-aggregation with α-synuclein. To test this, we performed tandem mass tag (TMT) quantitative proteomics on the detergent insoluble fractions from double mutant and α-syn mice. We found 185 proteins significantly increased in the double mutants (FDR-adjusted p < 0.05, Supplementary Table 4). Of these, 55 were ascribed to the “mitochondrion” Cell Component, which was significantly overrepresented (GO:0005739, FDR < 10^-15^). KEGG pathway analysis showed “oxidative phosphorylation” as the most enriched group of proteins that are increased in the insoluble fractions from double mutant spinal cords (Figure 6A-B, FDR < 10^-42^). This was not driven by increased total α-syn levels (Figure 6B). Interestingly, multiple neurodegenerative disease KEGG pathways were also enriched, most notably PD, suggesting clogging-induced protein aggregation may be clinically relevant in these conditions (Figure 6A). Finally, we detected a unique peptide within mitochondrial targeting presequence of COX5A, a protein significantly increased in the double mutant aggregates (Figure 6C). As this peptide is cleaved and degraded upon COX5A entry into mitochondria, this suggests the presence of un-imported mitochondrial preproteins within α-synuclein aggregates, which increases with import clogging. Taken together, the data suggest that mitochondrial protein import clogging may contribute to α-synuclein aggregation through co-aggregation with un-imported mitochondrial proteins.

**Figure 6.**
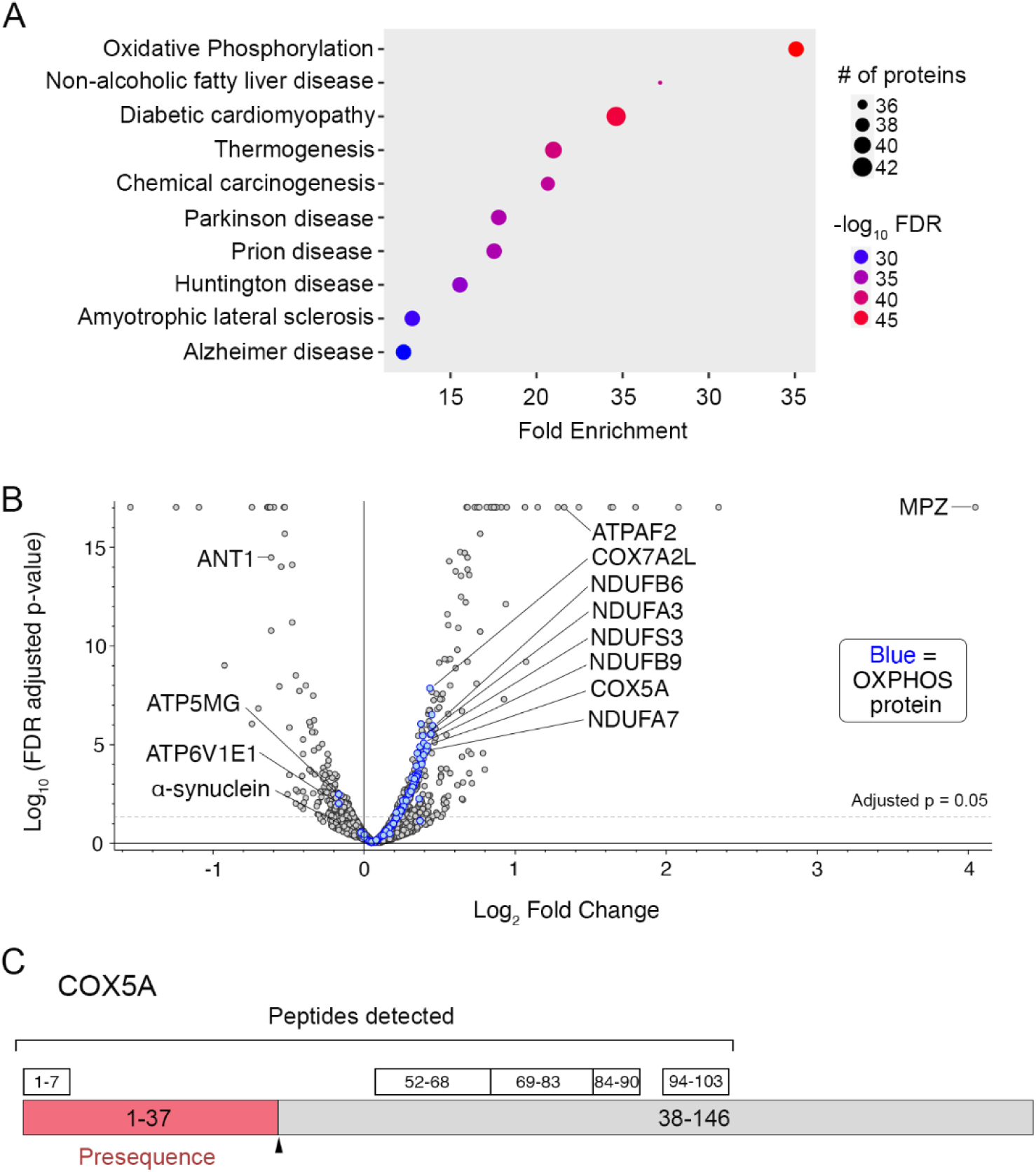
Mitochondrial protein import clogging increases co-aggregation of α-synuclein with mitochondrial preproteins *in vivo*. (A) KEGG pathway analysis of the proteins significantly increased (FDR adjusted *p* < 0.05) in the insoluble fraction of double mutant end-stage mouse spinal cord compared with α-syn alone, as determined by TMT-based quantitative proteomics. FDR, false discovery rate. (B) Volcano plot of TMT proteomics experiment with OXPHOS proteins labeled as blue dots. (C) Schematic of COX5A protein and the five peptides quantified in the TMT proteomics experiment, shown above the protein (38% coverage). In red is the COX5A pre-sequence (residues 1-37) that is normally cleaved and degraded if the protein is efficiently targeted into mitochondria.

## DISCUSSION

Two important hallmarks shared among prevalent neurodegenerative diseases such as Parkinson’s disease (PD), Lewy Body Dementia, Alzheimer’s disease, and ALS are mitochondrial dysfunction and cytosolic protein aggregation. Whether and how these hallmarks interact in disease pathogenesis is an open question. Studies in yeast and human cells over the last decade have shown that a diverse range of mitochondrial insults can cause the toxic accumulation and aggregation of mitochondrial preproteins in the cytosol, a process termed mitochondrial Precursor Overaccumulation Stress (mPOS) ^9,10^. The critical barrier to testing whether mPOS contributes to neurodegenerative diseases has been modeling protein import defects *in vivo* without confounding oxidative phosphorylation deficiency and/or lethal shutdown of mitochondrial biogenesis. In this study, we overcome this roadblock by establishing that mild protein import clogging by Ant1^p.A114P,A123D^ does not affect bioenergetics in the central nervous system of heterozygous mice. As a first step to assessing mPOS in neurodegeneration, we induced mitochondrial protein import clogging by expressing Ant1^p.A114P,A123D^ in an established mouse model of PD caused by expression of human α-synuclein with the A53T mutation, which is an autosomal dominant cause of familial PD and a highly aggregation-prone protein ^29^.

The data presented provide foundational evidence that reduced mitochondrial protein import efficiency can contribute to neurodegeneration involving α-synuclein aggregation independent of bioenergetics. Mechanistically, we show that mild import stress can increase α-synuclein aggregation and co-aggregation with un-imported mitochondrial preproteins, all of which correlates with a worsened neurodegenerative phenotype. We rigorously established that bioenergetic defects cannot explain the phenotype augmentation by Ant1^p.A114P,A123D^ in α-syn mice. These data suggest that the cytosolic accumulation of un-imported mitochondrial preproteins, i.e. mPOS, may indeed contribute to neurodegeneration independent of bioenergetics, adding a new layer to the complex relationship between mitochondrial dysfunction and neuropathogenic protein aggregates. This has broad pathophysiologic and therapeutic implications, and warrants testing of mPOS in further PD mouse models and other models of neurodegeneration such as Alzheimer’s and ALS.

We chose to test mPOS in a model of PD because it has long been appreciated that mitochondrial dysfunction can cause cytosolic α-synuclein aggregation, but the molecular mechanisms have not yet been elucidated. Indeed, toxins that inhibit mitochondrial complex I, such as MPTP and rotenone, cause α-synuclein aggregation and PD in humans, human cells, and animal models ^34–39^. Interestingly, MPTP can reduce mitochondrial protein import in cells as well as *in vitro* ^40^, and reduced protein import aggravates seeding of α-synuclein aggregates in cell models ^41^. Moreover, mitochondrial proteins are found in α-synuclein aggregates in PD patient brains ^42–45^. Thus, circumstantial evidence previously suggested that mPOS might contribute to α-synuclein aggregation. However, this is the first study to directly test this, and show that it occurs independent of bioenergetics.

Our finding that mild mitochondrial protein import defect is sufficient to alter the course of α-synuclein induced neurodegeneration could have important implications for better understanding of PD. As mentioned above, mitochondrial stress has been widely reported in PD patient samples. It is expected that genetic and environmental PD inducers (e.g., defective mitochondrial protein quality control and exposure to MPTP or rotenone) would reduce proton pumping across the IMM, which results in reduced membrane potential and protein import ^10^. On top of this, α-synuclein itself may impair mitochondrial protein import by binding the TOM20 import receptor ^46^ or through an effect on mitochondrial membrane potential ^47^. As expression and accumulation of α-synuclein increases in PD-affected neurons with normal aging ^48^, it is possible that an age-dependent increase in α-synuclein level gradually reduces mitochondrial protein import, thereby causing mPOS to aggravate its own aggregation in the cytosol. A similar process may happen in certain forms of familial PD, as loss of PINK1, which is a recessive cause of PD, has been convincingly shown to reduce mitochondrial protein import^49^.

In summary, we utilized our unique clogger mouse model to uncover the neurological effects specific to mitochondrial protein import stress and subsequent mPOS. We found that mild import stress can readily be mitigated in healthy mice but enhances protein aggregation and neurodegeneration in a mouse model of PD without affecting mitochondrial bioenergetics. Thus, mitochondrial protein import efficiency and mPOS may be disease modifying factors in neurodegeneration independent of OXPHOS.

## METHODS

### Mouse studies

All procedures were approved by the Animal Care and Use Committee (IACUC) at State University of New York Upstate Medical University and were in accordance with guidelines established by the National Institutes of Health. *Slc25a4* ^p.A114P,A123D^/+ mice, or “clogger” mice, were generate as previously described ^14^. After >10 back crosses with the C57BL/6NTac females (Taconic Catalog no: B6-F), male *Slc25a4* ^p.A114P,A123D^/+ mice were crossed with female C57BL/6 mice with transgenic expression of the human *SNCA* gene (coding for the α-synuclein protein) harboring the pathogenic A53T mutation (Jackson Lab #006823) ^29^. *SNCA* expression is driven by the mouse prion promoter in these mice. Where α-syn mice were tested, all wild-type and clogger control mice were littermates.

Animals in the α-syn background were examined daily for signs of illness for determination of “end stage” for lifespan, histological and biochemical analysis. An animal was considered “end stage” once it displayed paralysis and was so severely moribund that it was determined unlikely to survive more than an additional 48 hours, as judged by an experienced technician in our Department of Laboratory Animal Resources. A mouse was considered severely moribund if it also exhibited the following clinical signs: inability to eat, drink or eliminate; severe dehydration; labored breathing; severe lethargy. Other parameters used that were used in lifespan determination, but not specific to “end stage” α-syn mice were edema, sizable abdominal enlargement or ascites, significant skin lesions exposing muscle, progressive dermatitis, or a severely ulcerated or bleeding tumor.

### Mitochondrial isolation and purification

For mitochondrial respiratory studies, mice were sacrificed by decapitation without CO_2_ asphyxiation or anesthetic. Neural tissues were rapidly dissected (<60 seconds) and placed in 1 mL of ice-cold Isolation Medium (IM) (225 mM Mannitol, 75 mM sucrose, 5 mM HEPES-KOH pH 7.4, 1 mM EGTA, 0.1% BSA). Quick dissection of the spinal cord was enabled via hydraulic extrusion with ice-cold PBS ^50^. Tissue was minced on ice while submerged in IM as soon as possible. For crude mitochondrial fraction isolation (i.e. for cerebellum, spinal cord, and brain stem), well-minced tissue was homogenized with 4 strokes by hand in a 2 mL glass dounce homogenizer (pestle B, clearance 0.0005-0.0025 inches), followed by differential centrifugation at 4°C. First, homogenate was centrifuged at 2,000g for 5 minutes. Supernatant was then centrifuged at ∼21,000g for 20 minutes. Pellet was resuspended in 0.5 mL IM and centrifuged again at ∼21,000g for 20 minutes. Again, pellet was resuspended in 0.5 mL IM and centrifuged again at ∼21,000g for 20 minutes. The final pellet was resuspended in 0.2 mL IM.

Mitochondria were purified from the forebrain, which in our studies includes all neural tissue anterior to the junction between the cerebral cortex and the cerebellum. This was done using discontinuous Percoll gradient centrifugation, essentially as described ^51^. Minced forebrains were homogenized with three strokes with the pestle at 13,900 rpm in ice cold 7 mL IM. Homogenate was then centrifuged for 5 minutes at 2,000g. Resulting supernatant was centrifuged at 13,000 g for 12 minutes. Pellet was then resuspended in 2 mL IM containing 12% Percoll and used to build a discontinuous gradient using the following Percoll concentrations in isotonic IM: 5 mL 40%, 5 mL 23%, and ∼2.5 mL 12% (mito). Gradients were ultra-centrifuged in a swinging-bucket rotor (SW 41) at 18,500 rpm (∼43,000 g) for 20 minutes with slow acceleration and deceleration. Purified non-synaptosomal mitochondria were retrieved from the 23/40% interface, and synaptosomes were collected from the 23/12% interface. Both fractions were then diluted with ∼25 mL IM without Percoll, and centrifuged at 13,000g for 15 minutes. Mitochondrial pellet was resuspended in 1 mL IM and centrifuged at 13,000g for 12 minutes. Mitochondria were then washed two more times in 1 mL ice-cold IM, and finally resuspended in 80 μl for non-synpatosomal pure mitochondria, and 200 μl for synaptosomes. Protein concentration was then determined by Bradford assay.

### Mitochondrial oxygen consumption measurements

Oxygen consumption assays were performed using Oxygraph system form Hansatech, which has a small water-jacketed (37°C), magnetically stirred chamber sitting atop a Clark electrode. Each reaction occurred in 0.5 ml Respiratory buffer (125 mM KCl, 4 mM K_2_HPO_4_, 3 mM MgCl_2_, 1 mM EGTA, 20 mM HEPES-KOH pH 7.2). 150 μg or 300 μg of non-synaptosomal and synaptosomal mitochondria, respectively, were added first. Then, for Complex I-based respiration, reagents were added in the following sequence at the indicated final concentrations: Glutamate (5 mM) + Malate (2.5 mM); ADP (200 μM); oligomycin (3 μg/ml); FCCP (0.2 μM). For Complex II: rotenone (5 μM); succinate (10μM); ADP (300 μM); oligomycin (3 μg/ml); FCCP (0.2 μM).

### Cytochrome c oxidase activity assay

COX activity assay was performed essentially as described ^52^. Oxidized cytochrome c (Sigma-Aldrich: C2506) was reduced with sodium dithionate and dialyzed overnight in potassium phosphate buffer, followed by spectrophotometric confirmation that Cytochrome c was reduced (OD 550/560 ratio > 6). For the activity assay, snap-frozen crude spinal cord mitochondria were thawed on ice and 1 mL reaction tube was prepared with 20 mM potassium phosphate, pH 7.0, 0.45 mM DDM and 15 μM reduced cytochrome C. At time zero, 0.02 μg mitochondria were added to the reaction and OD550 was monitored every 2 seconds for 5 minutes. The slope over the first 12 seconds of the assay was used to determine the rate constants.

### Electron microscopy

For electron microscopy, *Slc25a4* ^p.A114P,A123D^/+ mice and littermate controls were processed as previously described ^53^. Briefly, mice were anesthetized with isoflurane and perfused intracardially with PBS initially, followed by fixative (1% paraformaldehyde, 1% glutaraldehyde, 0.12 M sodium cacodylate buffer pH 7.1, and 1 mM CaCl_2_). Perfused animals were refrigerated overnight, and CNS tissues dissected the next day and processed for TEM. The samples were examined with a JOEL JEM1400 transmission electron microscope and images were acquired with a Gaten DAT-832 Orius camera.

### Detergent solubility studies

Triton Lysis Buffer (0.5% Triton X-100, 150 mM NaCl, 50 mM HEPES-KOH pH 7.4, 1 mM EDTA with protease and phosphatase inhibitors (Roche)) was added to dissected and snap-frozen tissues at a ratio of 6 μL/mg of tissue. Tissue was minced and homogenized in a 2 mL dounce homogenizer, then incubated on ice for 30 minutes. Triton-insoluble fractions were collected by centrifugation at 21,000g for 20 minutes, then washed twice by vortexing in Triton Lysis Buffer. Soluble supernatant was centrifuged an additional two times at 21,000g for 30 minutes each. Triton insoluble fractions were solubilized with 5% SDS and 8M Urea, mixed with pipetting and sonicated for ∼10 seconds. Samples were then incubated at 42°C for 30 minutes before centrifugation at 16,000g for 20 minutes at room temperature. Supernatant was the final “Triton insoluble fraction” and was either directly loaded onto an SDS-PAGE gel for Western blot analysis or processed for quantitative mass spectrometry as described below.

### Sample processing for Quantitative Mass Spectrometry

Five biological replicates of α-syn and double mutant samples were prepared for multiplexed quantitative mass spectrometry as above. Samples were buffer exchanged on a 3 kDa molecular weight cutoff filter (Amicon 3k Ultracel) using 4 additions of 10 mM triethylammonium bicarbonate, pH 8.0 (Thermo). One hundred µg was taken for digestion using an EasyPep Mini MS sample prep kit (Thermo, A40006). To each buffer-exchanged sample, 65 µL of lysis buffer was added followed by 50 µL of reduction solution and 50 µL of alkylating solution. Samples were incubated at 95°C for 10 minutes, then cooled to room temperature. To each sample 5 µg of trypsin / Lys-C protease was added and the reaction was incubated at 37°C overnight. Half of each digest was used for subsequent labeling. TMT reagents were reconstituted with 40 µL acetonitrile (ACN) and the contents of each label added to a digested sample. After 60 min, 50 µL of quenching solution was added, consisting of 20% formic acid and 5% hydroxylamine (v/v) in water. The labeled digests were cleaned up by a solid-phase extraction device contained in the EasyPep kit, and dried by speed-vac. The individually labeled samples were dissolved in in 50 µL of 30% ACN and 0.2% trifluoroacetic acid (v/v) in water, and 15 µL of each was used to create a pooled sample consisting of 150 µg.

### Fractionation

Following an LC-MS experiment to check digestion and labeling quality of the pooled samples, 100 µg of the pooled sample was fractionated using a Pierce High pH Reversed-Phase Peptide Fractionation Kit (part # 84868), per the manufacturer’s instructions for TMT-labeled peptides. In brief, samples were dissolved in 300 µL of 0.1% trifluoroacetic acid in water and applied to the conditioned resin. Samples were washed first with water and then with 300 µL of 5% ACN, 0.1% triethylamine (TEA) in water. The second wash was collected for analysis. Peptides were step eluted from the resin using 300 µL of solvent consisting of 5 to 50% ACN with 0.1% TEA in eight steps. All collected fractions were dried in a speed-vac.

### LC-MS/MS

Dried fractions were reconstituted in 25 µL of load solvent consisting of 3% ACN and 0.5% formic acid in water, and a 5 µL aliquot was diluted 1:3 with the same solvent. Of these 15µL, 2 µL were injected onto a pulled tip nano-LC column (New Objective, FS360-75-10-N) with 100 µm inner diameter packed to 32 cm with 1.9 µm, 100 Å, C18AQ particles (Magic 2, Premier LCMS). The column was maintained at 50°C with a column oven (Sonation GmbH, PRSO-V2). The peptides were separated using a 135-minute gradient consisting of 3 – 12.5% ACN over 60 min, 12.5 – 28% over 60 min, 28 - 85 % ACN over 7 min, a 3 min hold, and 5 min re-equilibration at 3% ACN. The column was connected inline with an Orbitrap Lumos (Thermo) via a nanoelectrospray source operating at 2.3 kV. The mass spectrometer was operated in data-dependent top speed mode with a cycle time of 3s. MS^1^ scans were collected from 375 – 1500 m/z at 120,000 resolution and a maximum injection time of 50 ms. HCD fragmentation at 40% collision energy was used followed by MS^2^ scans in the Orbitrap at 50,000 resolution with a 105 ms maximum injection time.

### Database searching and reporter ion-based quantification

The MS data was searched using SequestHT in Proteome Discoverer (version 2.4, Thermo Scientific) against the *M. Musculus* proteome from Uniprot, containing 50961 sequences and a list of common laboratory contaminant proteins. Enzyme specificity for fully tryptic with up to 2 missed cleavages. Precursor and product ion mass tolerances were 10 ppm and 0.02 Da, respectively. Cysteine carbamidomethylation, TMT 10-plex at any N-terminus and TMT 10-plex at lysine were set as a fixed modifications. Methionine oxidation was set as a variable modification. The output was filtered using the Percolator algorithm with strict FDR set to 0.01. Quantification parameters included the allowance of unique and razor peptides, reporter abundance based on intensity, lot-specific isotopic purity correction factors, normalization based on total peptide amount, protein ratio based on protein abundance, and background-based hypothesis testing (t-test). Final protein list was analyzed for enrichment using STRING database version 11.5 ^54^ and ShinyGO 0.77 ^55^.

### Immunoblot analysis

Purified brain mitochondria were solubilized in Laemmli buffer and processed for Western blotting using standard procedures and Total OXPHOS Antibody Cocktail (#ab110411, Abcam).

### Mouse perfusion and lumbar spine immunofluorescence imaging

Mice were perfused via the heart as described previously ^56^. Briefly, animals were anesthetized with inhaled isoflurane prior to sequential cardiac perfusion with PBS (pH 7.4) followed by fresh 4% PFA dissolved in PBS (pH 7.4). The brain and spinal cord were removed, cryoprotected in increasing concentrations of sucrose and embedded in OCT. Cryosections of the lumbar spinal cord were obtained at 30mm, placed on Superfrost plus slides (FisherScientific #12-550-15), and stored airtight at −70°C prior to immunofluorescence staining. Slides were thawed, washed in PBS containing .05% Tween 20 and incubated for one hour in blocking buffer (10% normal goat serum + .05% Tween 20 +.3% Triton-x-100 dissolved in PBS). Blocking buffer was discarded and slides were washed and incubated with primary antibody: rabbit anti-Alpha-synuclein (phospho S129) (Abcam #ab51253, 1:10000) diluted in blocking buffer overnight. Primary antibody was discarded, and slides were washed and incubated with secondary antibody: Alexa Fluor 633 conjugated goat anti-rabbit IgG (H+L) (Invitrogen #A-21070, 1:500) diluted in blocking buffer for 2 hours. Secondary antibody was discarded, and slides were washed and mounted with ProLong gold antifade mountant with DAPI (Invitrogen #P36391). Z stack confocal images of the lumbar spine anterior horn were obtained using the Leica SP8 confocal microscope. For image analysis, raw Leica image files were imported into FIJI ^57^and the brightest z plane for each section was quantified.

### Central nervous system dissection

Neural tissues were dissected and snap-frozen as quickly as possible, and the spinal cord ejected from the spinal canal via hydraulic extrusion with ice cold DEPC-treated PBS. Accurate identification of the striatum was less straight forward than the cerebellum and spinal cord. First, cerebellar, brain stem, and olfactory bulb were removed from the forebrain. The brain was then cut sagittally along the interhemispheric fissure. Approaching from the medial aspect of the brain, inner brain material (including the thalamus, septum and underlying striatum) was removed by cutting just below the corpus collosum, thus separating these tissues from the hippocampus and associated cerebral cortex. The easily identifiable thalamus and hypothalamus were removed, and the remaining tissue from this “inner brain” material was labeled striatum. Tissue was frozen in liquid nitrogen.

### RNA purification and sequencing

Tissue was disputed in QIAzol lysis reagent, using the Qiagen TissueRuptor. RNA was then extracted using the Qiagen miRNeasy Mini Kit. RNA quality and quantity were assessed with the RNA 6000 Nano kit on the Agilent 2100 Bioanalyzer. For striatum, sequencing libraries were prepared using the Illumina TruSeq Stranded mRNA Library Prep kit, using 1ug total RNA as input. For cerebellum and spinal cord, sequencing libraries were prepared using the Illumina Stranded mRNA Library Prep kit Ligation, using 500ng Total RNA as input. The two library prep kits are equivalent. Library size was assessed with the DNA 1000 Kit on the Agilent 2100 Bioanalyzer. Libraries were quantified using the Quant-IT High Sensitivity dsDNA Assay (Invitrogen) on a Qubit 3.0 Fluorometer. Libraries were sequenced on the NextSeq 500 instrument, with paired end 2×75bp reads.

### RNAseq analysis

All analysis was performed on Partek Flow Genomic Analysis Software. All reads were aligned to the *mus musculus* genome (mm10) using STAR – 2.7.3a and quantified using Partek E/M. Reads were normalized with TPM (transcripts per million) and quantile normalization including all samples from spinal cord, cerebellum and striatum. We probed for differentially expressed genes using ANOVA, analyzing data from all tissues together. Within-tissue analyses shown as heat maps, volcano plots, and gene lists were generated in Partek from this analysis. Similarly, pathway analysis shown for the striatum data was performed using Partek Pathway Analysis software. For volcano plots, *Slc25a4* transcript was omitted to facilitate visualization.

### Behavioral Assays

Behavioral assays were performed in a particular order to minimize the likelihood that one test affects mouse behavior on subsequent days. The order went as follows: elevated plus maze, Y-maze spontaneous alternation test, open field activity test, novel object recognition test, Morris Water Maze, puzzle box test, followed by rotarod testing. Male and female mice were tested in all assays. Each data point shown in behavioral assays represents an independent mouse. Elevated plus maze, y-maze spontaneous alternation, and open field activity testing were done on two independent cohorts. For every test, mice were habituated to the testing room for 30 minutes prior to each test, and odors and residue was removed after each test with 70% ethanol. Mouse activity and scoring in each test was automatically measured using ANY-Maze behavioral tracking software, except for puzzle box, rotarod and beam walking tests, which were scored manually with the experimenter blinded to genotype. Elevated plus maze, y-maze, open field activity, and novel object recognition were performed as previously described ^58^.

### Elevated plus maze (EPM)

EPM is used to assess anxiety-like behavior. We used a standard mouse EPM apparatus from San Diego Instruments, which consists of two closed arms and two open arms perpendicular to one another, forming a “plus” shape. Mice were placed in the center of the apparatus, facing an open arm, and allowed to explore freely for 5 minutes under ambient light. The time spent in the open arms is reported to reflect anxiety-like behavior, such that the less time in the open arms, the more anxiety-like the behavior is.

### Y-maze Spontaneous Alternation Test

We used the Y-maze Spontaneous Alternation Test to assess spatial working memory. A custom-built apparatus was used, which consisted of 3 walled arms 16 inches long that are angled 120° from one another. Mice were placed in the center and allowed to freely explore for 5 minutes in dim light. This test is based on the rodent’s tendency to explore new environments. Exploring the 3 arms consecutively, without re-entry into an arm, is called a triad. This implies that the mouse “remembers” which arm it was most recently in, despite all arms appearing identical. To control for difference in total number of arm entries, we report Fraction of Alternation, which is (total number of triads) / (total number of arm entries – 2). Mice with less than 5 total arm entries were excluded from Fraction of Alternation analysis.

### Open field activity test

Open field activity testing was performed to assess spontaneous locomotor activity and anxiety-like behavior. We used a standard apparatus from San Diego Instruments, which consists of 50 cm x 50 cm open field surrounded by non-transparent walls. Mice were allowed to explore freely for 10 minutes in ambient light. The total time spent away from the pre-designated “center zone” is reported to reflect anxiety-like behavior.

### Novel object recognition (NOR) test

NOR testing was performed to assess long-term object recognition memory and was performed in the open field apparatus. Briefly, two identical objects (cubes) were placed in the chamber and the mice were allowed to explore the objects for 5 minutes per day in dim lighting for two training days. On the third day, a novel object (cylindrical piece of wood) replaced one of the cubes, and mice were allowed to explore the chamber for 10 minutes. If the mice remember the objects from the training days, then they tend to spend more time exploring the novel object. Reported is the time spent interacting with the novel object on the final day of testing. Mice with less than 5 seconds of total object exploration time on the final testing day were excluded from discrimination index analysis.

### Morris water maze

Morris water maze was performed to assess long-term spatial memory and learning, and was performed in an inflatable hot tub with the water temperature at 26°C in dim lighting, essentially as previously described ^59^. Briefly, mice were placed in the circular pool for 4 trials per day and allowed one minute to find a platform that they can stand on. Platform was made of transparent plastic and placed just beneath the water surface as to make it invisible to the rodents. For pre-training day 1, mice were placed on the platform for one minute, followed by placement of the mouse proximally to the platform so it learns it can escape the water. On pre-training day 2, mice were placed in a pseudo-randomized quadrant of the pool and allowed one minute to escape to the platform, which was made visible on this day with a small flag. Then, for 7 training days, the platform was below the surface with no flag, and the mice were given 4 trials per day to learn where the platform was using spatial cues from around the room. On the probe day, the day after training day 7, the platform was removed, and the amount of time the mice spent swimming in the quadrant of the pool that previously had the platform, as well as the number of times the mice entered the platform zone, were recorded. The better long-term spatial memory the rodent has, the more time it spends in the target quadrant, and the more times it will enter the platform zone.

### Puzzle Box Test

To assess executive function, we performed the puzzle box test as previously described ^19, 20^. Briefly, mice were placed in a custom-built 60 cm x 28 cm apparatus that is well-lit with a small fan blowing into it to create a stressful environment with multisensory stimuli. The mice are given a passage (5 cm by 5 cm doorway) to small and dark target location filled with home bedding to make the mice comfortable. The idea is to test the mouse’s ability to “problem solve” by removing obstacles between the stressful environment and the target location. In condition 0, there is no obstacle. Condition 1 had a U-shaped channel within the doorway. Condition 2 had the same U-shaped channel within the doorway, but with a mound of cage bedding placed in it to block the doorway. Condition 3 had a crumped piece of paper placed in the doorway. Finally, condition 4 had a cube placed in the doorway, with a raised edge on top of it such that the mice had to pull it out of the doorway and could not push it through. We manually recorded the time it took to reach the target location and plotted that as “escape latency”. A maximum of 5 minutes was allowed for each trial.

### Beam walking test

Beam walking test was used as a measure of balance and coordination. Although typically the primary read-out for this assay is time to traverse the beam, analysis of these data was compromised by the substantially increased locomotor activity of transgenic α-synuclein^A53T^ mice. Thus, we prioritized the number of times each mouse’s hind limbs slipped off the beam as the primary read-out for balance and coordination.

We custom-built a beam walking apparatus, consisting of a one-meter-long wooden beam suspended ∼16 inches above the floor. The start end of the beam was suspended in the air without an escape route, and the other end has the target box for the mice to escape into. The target box is an enclosed black cube of about 9 inches on each side. Before each test, home bedding from the tested mouse is added to the escape box for enticement. Aversive stimuli (a bright LED light and a small fan) were placed at the start end of the beam such that the escape box was the only route to protection. Mice were trained on an easy beam (0.5-inch rectangle) over two days. In training, if a mouse stopped moving while on the beam it was gently prodded by the experimenter. On training day 1, the mouse was placed directly in the target box for one minute, then placed on the beam two inches from the target box and allowed to escape to the target box, and finally placed at the far end of the beam for full traverse of the beam. On training day 2, each mouse was given 3 more trials on the easy beam. On test day, each mouse was tested twice on 0.5-inch circular and 0.25-inch rectangular beams. The number of hindlimb slips per beam was manually scored by an experimenter who was blinded to genotype. Plotted in Figure 4 is the average number of slips of the two trials.

### Accelerating rotarod test

Mice were initially acclimated to the rotarod for 30 seconds at 4 rotations per minute (rpm). Mice aged 19-22 months (n > 16 mice per genotype, n > 5 per sex per genotype) were tested on a high difficulty rotarod protocol of acceleration from 4 to 40 rpm over 2 minutes, three trials per day for two consecutive days. The time at which the mouse fell off the rod was recorded and plotted with the experimenter blinded to genotype.

### Treadmill exhaustion test

Treadmill tests were performed on an Exer 3/6 animal treadmill at a 5° incline (Columbus Instruments). For one week, mice were familiarized with the treadmill with 10-minute running sessions at 10 m/min every other day. For resistance testing, the treadmill started at 9 m/min with 1.2 m/min/min acceleration until the mice reached exhaustion. For power testing, the treadmill started at 9 m/min with 1.8 m/min/min acceleration until the mice reached exhaustion. Mice were deemed exhausted when they remained in continuous contact with the shock grid for 5 seconds. Three full days of rest were allowed between training and each testing day.

### Optomotor response test

Optomotor acuity and contrast sensitivity of mice were determined by observing their optomotor behavior responses to a rotating sine wave grating stimulus using the OptoMotry© system ^60^ as described previously ^61, 62^. Briefly, mice were placed on a pedestal at a center of the OptoMotry chamber enclosed by four computer monitors. The computer randomly displayed sinusoidal pattern gratings rotating in a clockwise or counter-clockwise direction. The observer was blind to the direction of rotation and chose the direction of pattern rotation based on the animal’s behavior. Auditory feedback indicated to the observer whether the selected direction was correct or incorrect. Using a staircase paradigm, the computer program controlled the spatial frequency and contrast of the stimulus. The threshold was set at 70% correct responses. Contrast sensitivity was defined as the reciprocal of the threshold contrast value and was measured at 1.5 Hz and a spatial frequency of 0.128 cycles/degree. Acuity was measured at a speed of rotation of 12 degrees/s and 100% contrast. All measurements were performed at the unattenuated maximal luminance of the OptoMotry© system (∼70 cd/m^2^, producing approximately 1500 R*/rod/s ^63^).

### Statistical Analysis

Statistical analyses were performed using GraphPad Prism. For behavioral tests, we always probed for a statistically significant effect of Genotype, Sex and Genotype x Sex interactions. If data are presented without separating sex, this indicates there was no significant main effect of sex and no significant Genotype x Sex interaction. For details on statistical testing of specific data, please see Figure Legends.

## ACKNOWLEDGEMENTS

We thank Ebbing de Jong and Auyon J. Ghosh for help in the analysis of proteomic data. This work was supported by National Institutes of Health pre-doctoral fellowship F30AG-060702 to L.P.C., American Heart Association pre-doctoral fellowship # 906215 to A.R., and National Institute of Health grants R01AG063499 (to X.J.C. and F.M.) and R01AG061204 (to X.J.C.).

## AUTHOR CONTRIBUTIONS

L.P.C. and X.J.C. conceived experiments and wrote the manuscript. L.P.C., A.R., F.M. and X.J.C. secured funding. L.P.C., A.R., X.W., S.B., and Y.U. performed experiments. F.M. provided expertise in designing and performing moue behavioral assays and RNA-seq analysis. E.C.S. participated in data analysis.

## MATERIALS AND CORRESPONDENCE

Material requests and correspondence should be addressed to X.J.C..

## SUPPLEMENTAL INFIRMATION

**Figure S1.**
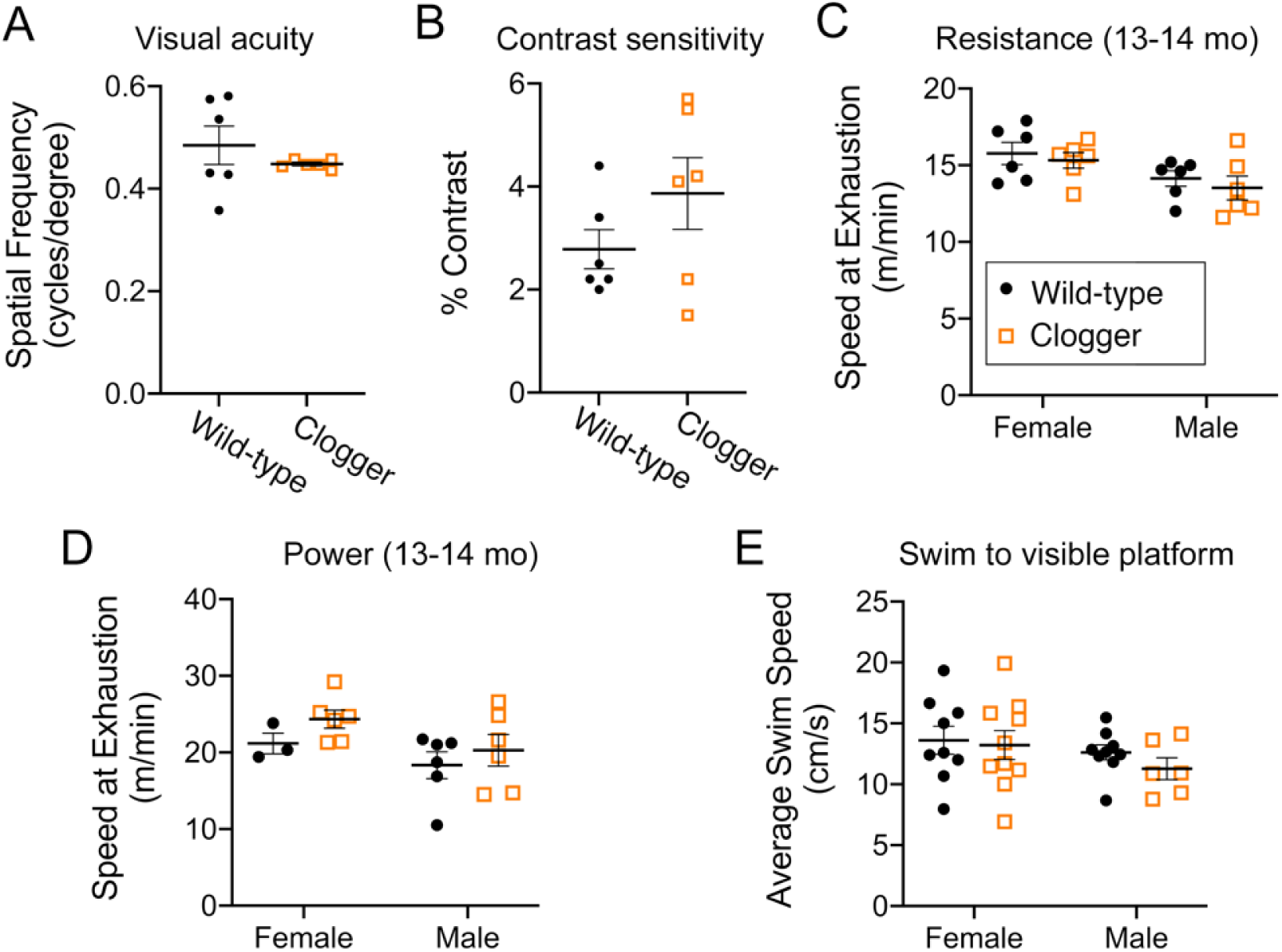
Visual and skeletal muscle functions are not impaired in middle aged clogger mice. (A) Visual acuity of clogger mice is intact at 17-18 months of age, as suggested by optomotor response testing (n=3/sex/genotype). (B) Visual contrast sensitivity of clogger mice is intact at 17-18 months of age, as suggested by optomotor response testing (n=3/sex/genotype). (C) Clogger mice do not show exercise intolerance at 13-14 months of age, as suggested by similar time to exhaustion when forced to run on a slowly accelerating treadmill (“resistance test”). (D) Clogger mice do not show exercise intolerance at 13-14 months of age, as suggested by similar time to exhaustion when forced to run on a quickly accelerating treadmill (“power test”). (E) Clogger mice swim to a visible escape platform at similar average speeds as wild-type mice, suggesting preserved muscle function. These mice were 19-22 months of age.

**Figure S2.**
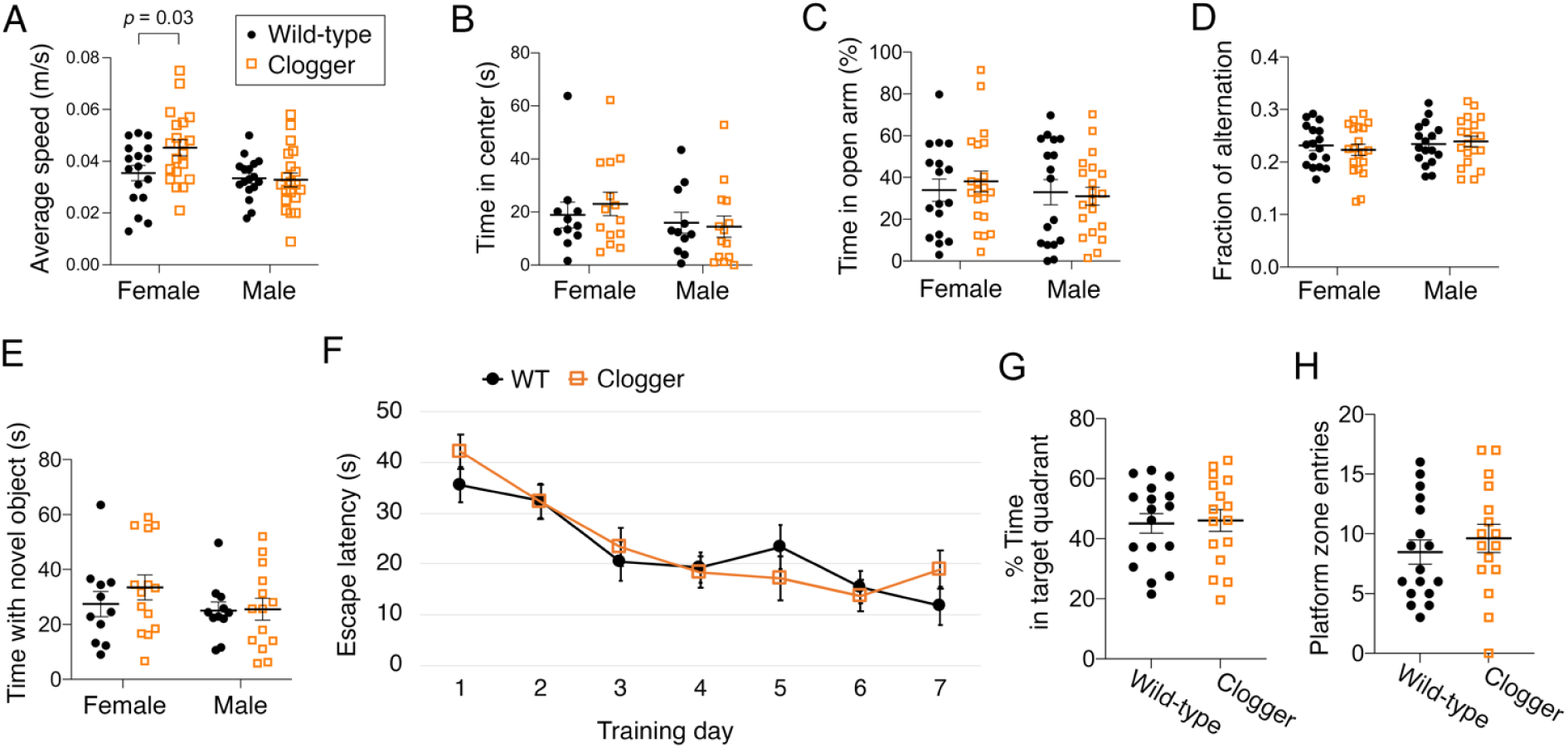
Clogger mice demonstrate sex-dependent increases in locomotor activity, but no changes in anxiety-like behavior or memory. (A) Female clogger mice have increased spontaneous locomotor activity. Two independent aged mouse cohorts (14-17 months of age) were monitored in the open field apparatus and average speed throughout the 10 minutes was plotted. Data were analyzed with a two-way ANOVA with Sidak’s multiple comparison’s test. Each dot is a unique mouse. (B) Clogger mice show unchanged levels of anxiety-like behavior, as indicated by the amount of time mice spent in the center vs the edges of the open field apparatus. (C) Clogger mice show unchanged levels of anxiety-like behavior, as indicated by the amount of time mice spent in the open vs closed arms in the elevated plus maze. (D) Clogger mice do not have impaired spatial working memory, as indicated by Y-maze testing. Fraction of alternation = (# of complete triads) / (# arm entries – 2). A triad is when a mouse enters all three arms consecutively without re-entering a previously visited arm in between. (E) Clogger mice do not have impaired long-term object memory, as indicated by the amount of time mice spent interacting with a novel object. (F) Clogger mice do not have spatial learning defects, as indicated by Morris Water Maze testing. Plotted is latency to escape from water to a hidden platform during 7 training days (n=4 trials/day). (G) Clogger mice do not have impaired long-term memory, as indicated by the probe trial after 7 days of training in the Morris Water Maze. The target quadrant of the swimming pool contained the hidden platform during training but was removed for the probe trial. (H) Clogger mice do not have impaired long-term memory, as indicated by the probe trial after 7 days of training in the Morris Water Maze. Plotted is the number of times the mice swam into the “platform zone”, i.e. the area where the hidden platform was during training days. N = 16 or 17 mice/genotype, > 5 mice/sex/genotype for F-H.

**Figure S3.**
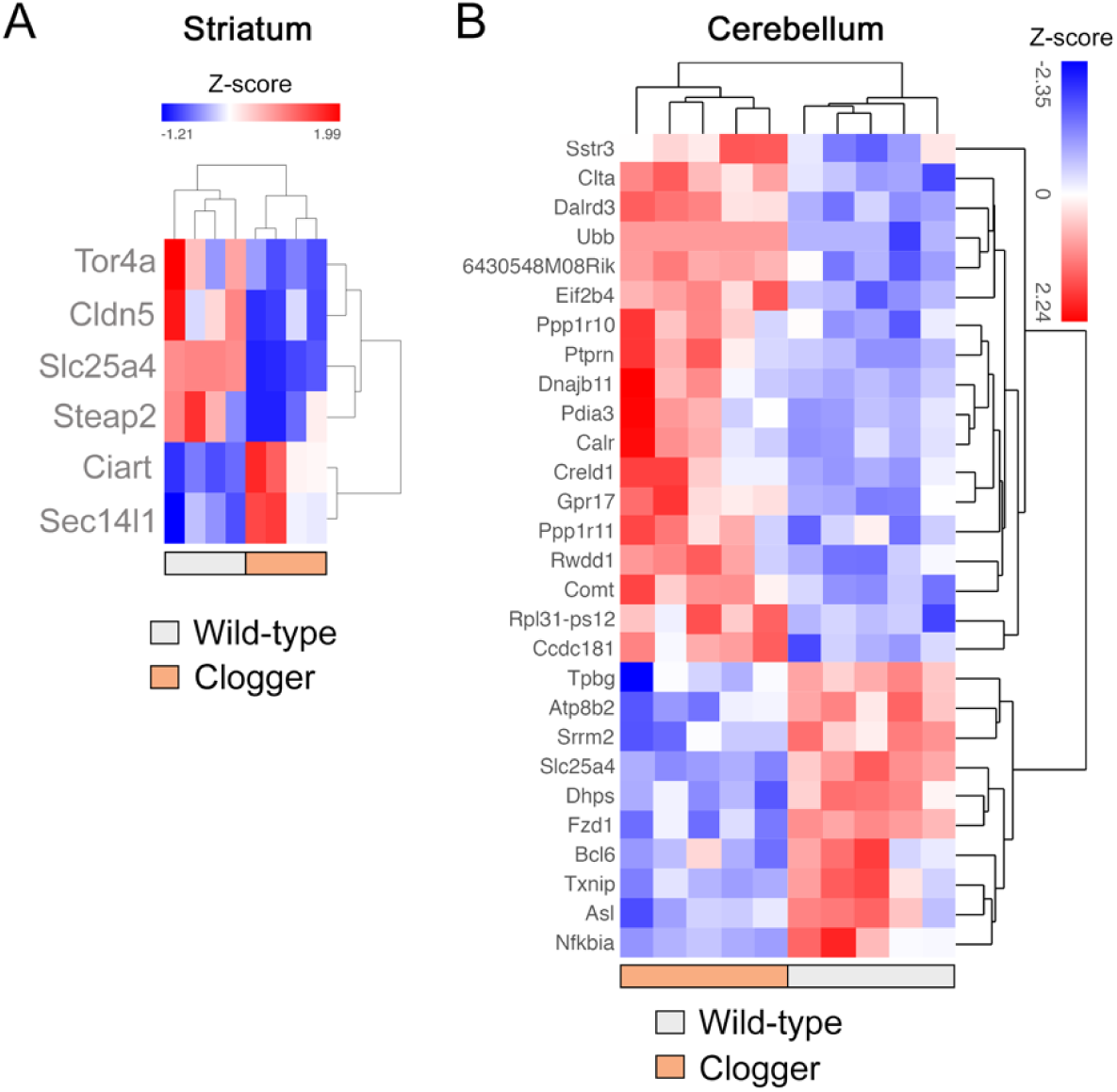
Transcriptomes of the striatum and cerebellum of Clogger mice reveal minimal changes. Note that *Slc25a4* (*ANT1*) transcript level is consistently lower due to reduced transcription or partial instability of the mutant *Ant1^p.A114P, A123D^* allele (our unpublished observation). (A) Hierarchical clustering heatmap of all genes that are differentially expressed in the striatum of clogger mice, compared with wild-type (*q*<0.05). (B) Heatmap of all genes that are differentially expressed in the cerebellum of clogger mice, compared with wild-type (*q*<0.05).

**Figure S4.**
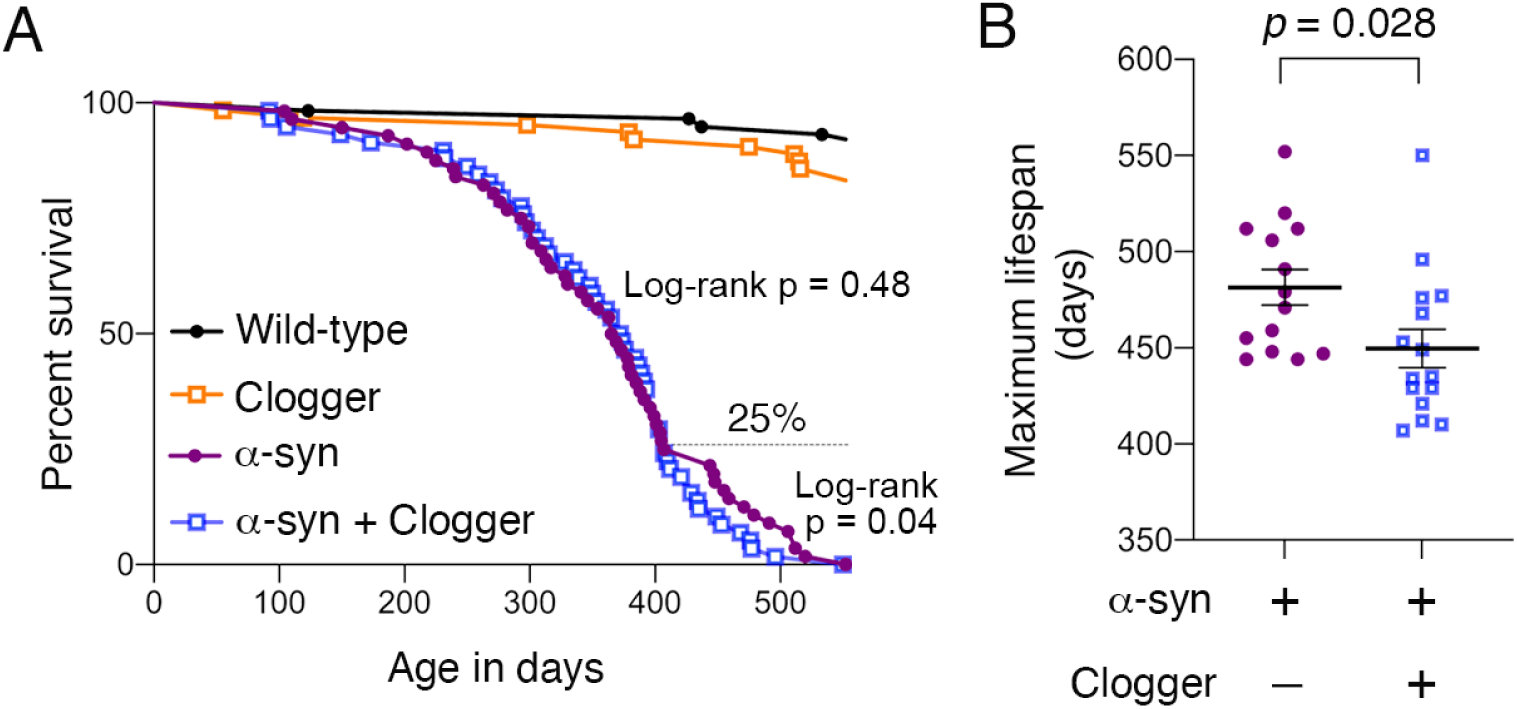
Lifespan analyses of wild-type, Clogger, α-syn, and double mutant mice. (A) Lifespan analysis of at least 56 mice per genotype. The 25% longest-lived mice depicted in (E) are noted on the graph. (B) Maximum lifespan analysis of the 25% longest-lived mice suggests modestly reduced maximal lifespan in double mutant mice. Data were analyzed by unpaired student’s *t* test.

**Figure S5.**
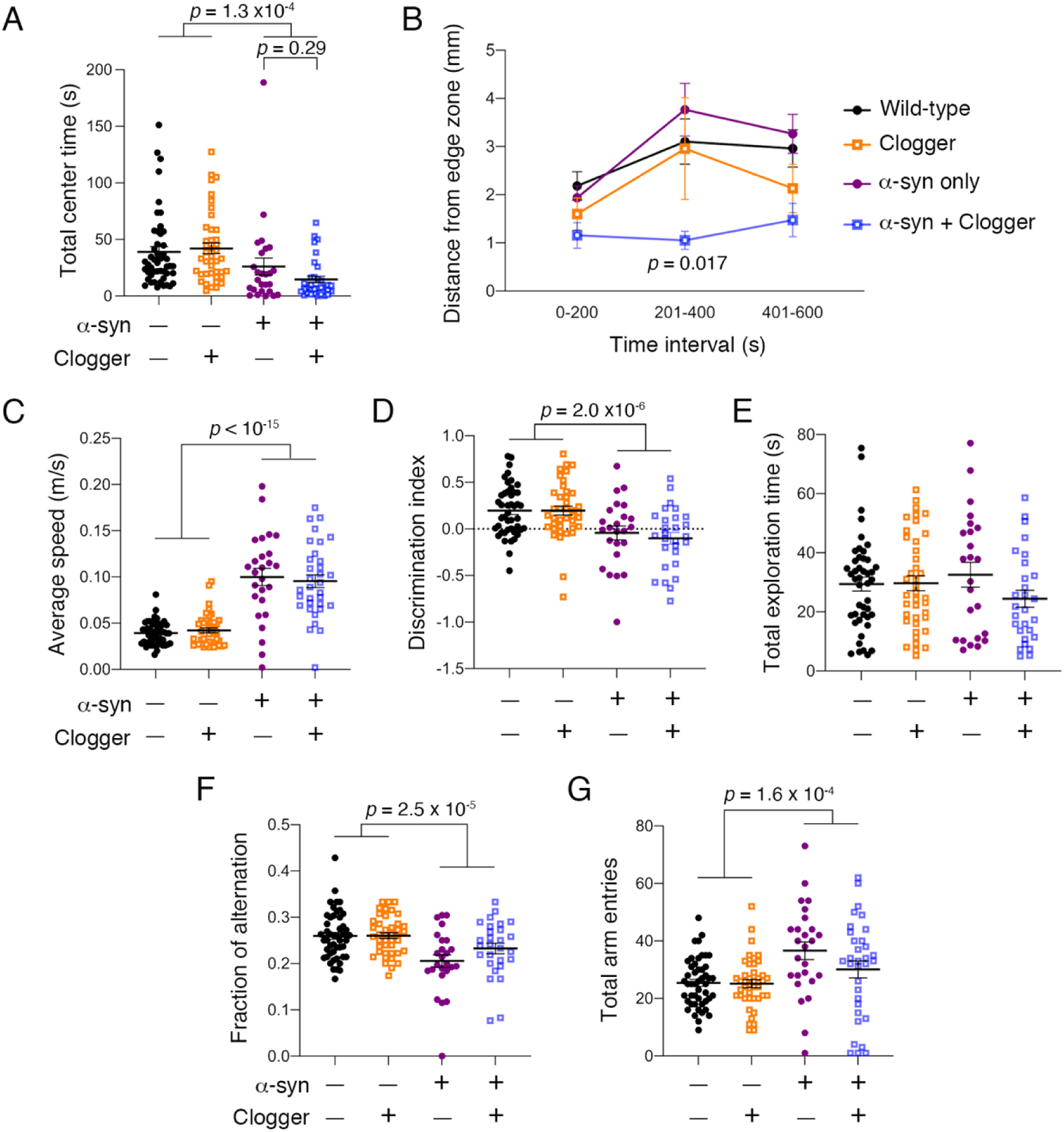
Assessment of anxiety-like behavior, spontaneous locomotion, and memory in wild-type, Clogger, α-syn, and double mutant mice. (A) α-synuclein^A53T^ expression significantly increases anxiety-like behavior, as indicated by spending less time in the center zone in the open field test. While clogger protein expression reduced the average time spent in the center zone compared in α-syn transgenics, this was not statistically significant. (B) Open field test data were divided into 200-second epochs, and the average distance from the edge zone while in the center zone was plotted. This showed that double mutant mice tend to stay closest to the edge zone, particularly in the middle epoch. Data were analyzed using two-way repeated measures ANOVA with epoch as the within-subjects variable and a-syn and clogger genotypes as the between-subjects variables, with Geisser-Greenhouse correction for violation of sphericity. A significant interaction effect was detected (Epoch x a-syn x clogger, *p* = 0.042). Dunnett’s multiple comparisons test detected a significant reduction in average distance from the edge zone for double mutant mice in the middle epoch (*p* = 0.017, shown as *). (C) α-synuclein^A53T^ expression significantly increases average speed of spontaneous locomotion in the open field test, which was not affected by the clogger protein. (D) α-synuclein^A53T^ expression significantly impairs long-term object memory in the novel object recognition test, which was not affected by the clogger protein. For better visualization, the discrimination index was plotted, which = ((time with new object) – (time with old object)) / (total object exploration time). (E) α-synuclein^A53T^ expression nor clogger protein expression affected total object exploration time on the probe trial of the novel object recognition test. (F) α-synuclein^A53T^ expression significantly impairs working spatial memory, as suggested by reduced fraction of alternation in the y-maze. This effect was not significantly affected by clogger protein expression. (G) a-synuclein^A53T^ expression significantly increases the total number of arm entries in the Y-maze spontaneous alternation test, which is not affected by clogger protein expression. All data in this figure were generated from 8-9 month old mice. Data were first analyzed with three-way ANOVAs probing for effects of sex, α-synuclein^A53T^, and clogger genotype. With no significant effect or interaction effect from sex, males and females were consolidated, and two-way ANOVAs were performed to simplify post-hoc analyses. For (A), (C), (D), (F), and (G), two-way ANOVA showed a significant main effect of α-synuclien^A53T^ expression, as indicated.

**Figure S6.**
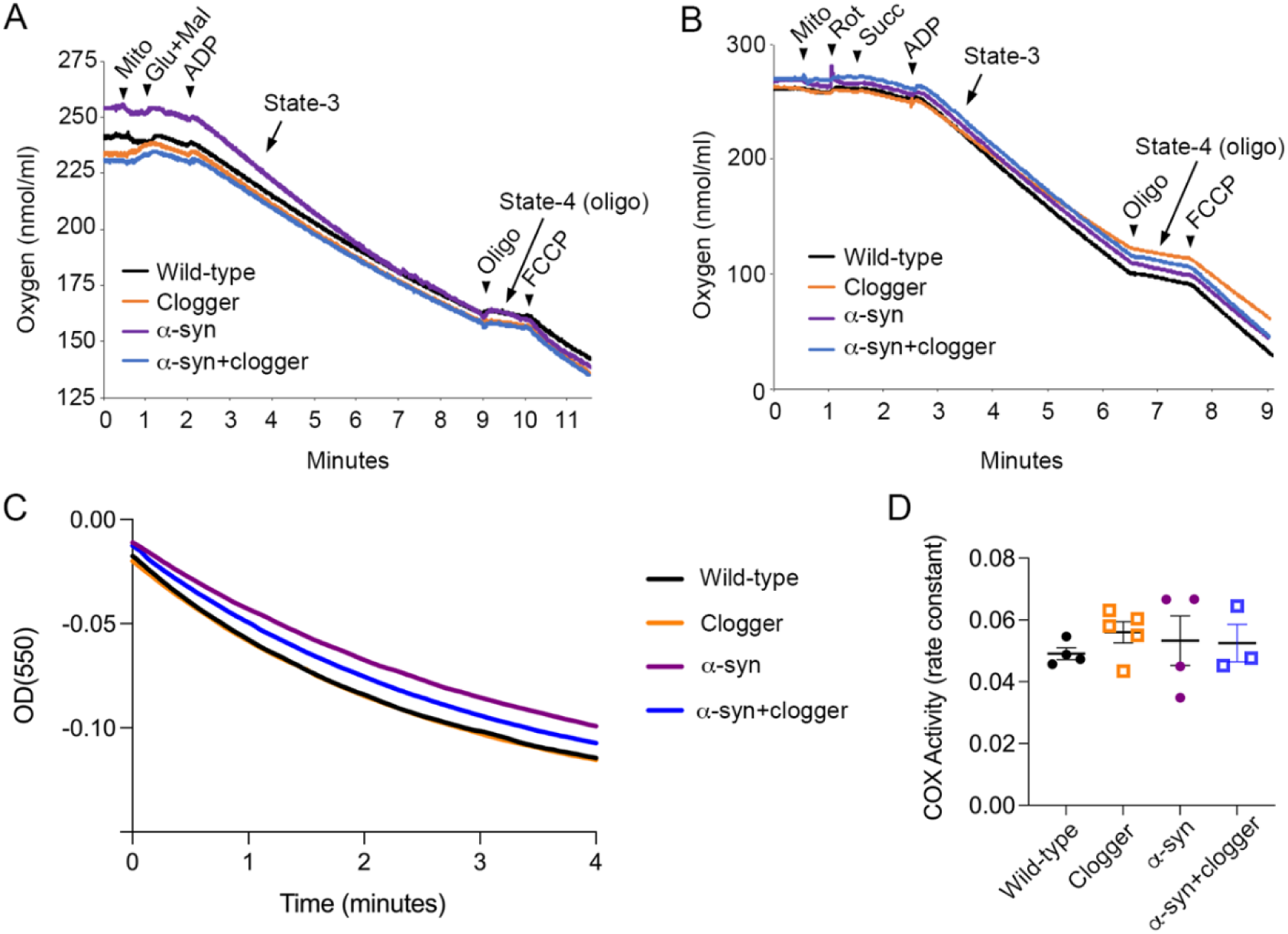
No mitochondrial bioenergetic defects found in synaptosomal fractions of Clogger, α-syn, and double mutant mice. (A) Oxygen traces from respiration measurements of brain synaptosomal fractions was measured after stimulation with complex I stimulated by glutamate (glu) and malate (mal). FCCP, Trifluoromethoxy carbonylcyanide phenylhydrazone; Oligo, oligomycin; Glu, glutamate; Mal, malate. State 3 respiration is the maximal respiratory rate after addition of ADP. State 4 respiration is the respiratory rate after depletion of ADP. (B) As in (A), except with complex I inhibited with rotenone and complex II stimulated with succinate. (C) Spectrophotometric traces from assessment of complex IV activity from crude mitochondrial fractions isolated from 9 months old mouse spinal cords. (D) Quantitation of biological replicates of the Complex IV activity rate constants determined from (C). Each dot represents an independent biological replicate.

**Figure S7.**
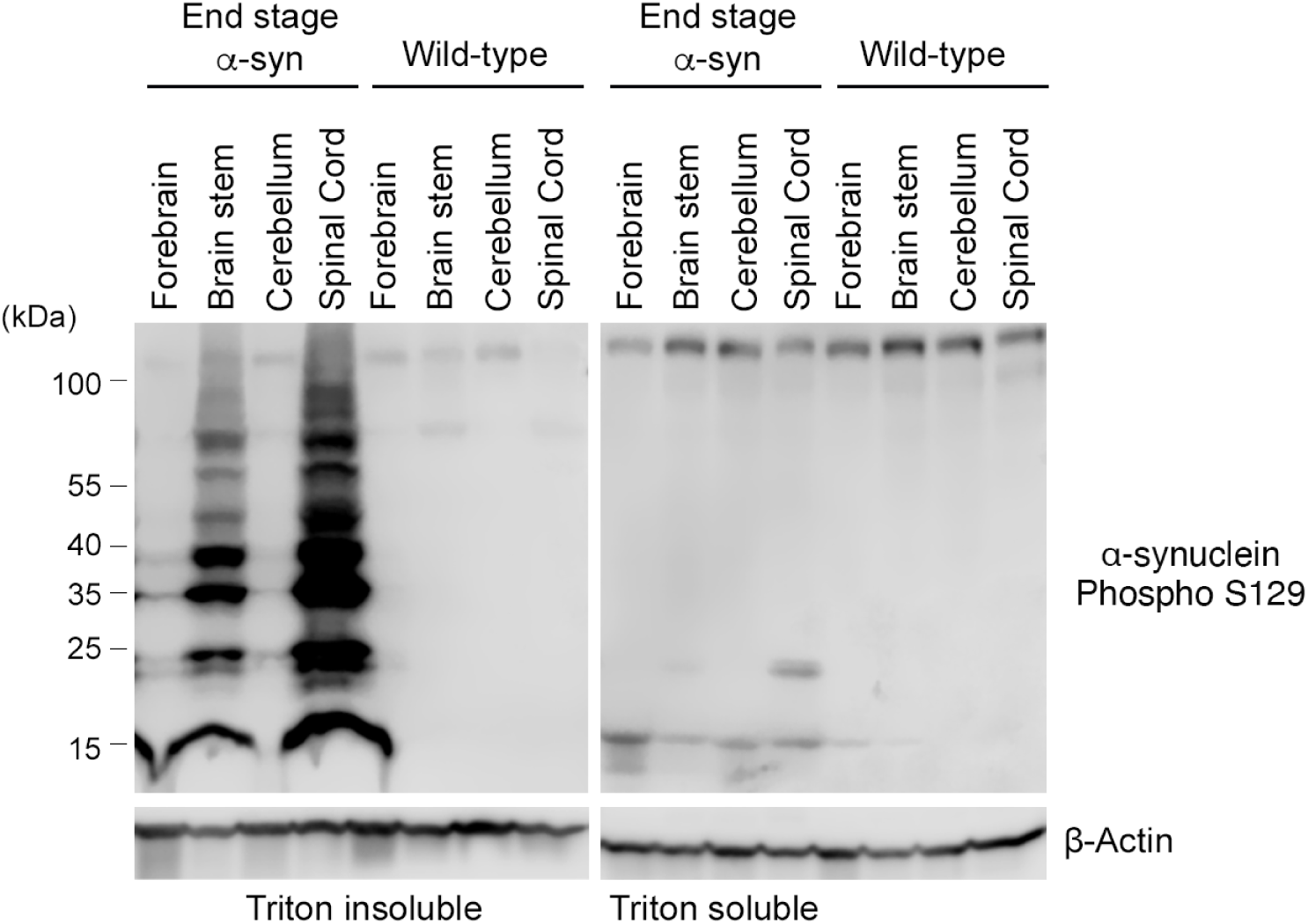
Phosphorylated α-synuclein is preferentially insoluble in the spinal cord of end-stage transgenic α-synuclein(A53T) mice. Immunoblot analysis of central nervous system tissues solubilized with 0.5% triton to assess the tissue distribution of aggregated α-synuclein. Antibody is specific to α-synuclein phosphorylated at Ser129.

### SUPPLEMENTAL EXCEL TABLES

**Supplemental Table 1** – RNA-seq data of spinal cord from *Ant1^p.A114P, A123D^*/+ compared with wild-type mice.

**Supplemental Table 2** – RNA-seq data of cerebellum from *Ant1^p.A114P, A123D^*/+ compared with wild-type mice.

**Supplemental Table 3** – RNA-seq data of striatum from *Ant1^p.A114P, A123D^*/+ compared with wild-type mice.

**Supplemental Table 4** – TMT mass spectrometry analysis of spinal cord Triton-insoluble proteome from *Ant1^p.A114P, A123D^*/+ α-syn double mutant compared with α-syn single mutant mice.

